# Controlling evolutionary dynamics to optimize microbial bioremediation

**DOI:** 10.1101/705749

**Authors:** Shota Shibasaki, Sara Mitri

**Author notes:** Author contribution: SS and SM conceived of the project, SS built and analyzed the model, SS and SM wrote the manuscript. E-mail addresses: SS, SM. Corresponding author’s detail. Data availability: Python code and csv files used for simulations and analytic calculations are available at GitHub.

## Abstract

Some microbes have a fascinating ability to degrade compounds that are toxic for humans in a process called bioremediation. Although these traits help microbes survive the toxins, carrying them can be costly if the benefit of detoxification is shared by all surrounding microbes, whether they detoxify or not. Detoxification can thereby be seen as a public goods game, where non-degrading mutants can sweep through the population and collapse bioremediation. Here, we constructed an evolutionary game theoretical model to optimize bioremediation in a chemostat initially containing “cooperating” (detoxifying) microbes. We consider two types of mutants: “cheaters” that do not detoxify, and mutants that become resistant to the toxin through private mechanisms that do not benefit others. By manipulating the concentration and flow rate of a toxin into the chemostat, we identified conditions where cooperators can exclude cheaters that differ in their private resistance. However, eventually, cheaters are bound to invade. To overcome this inevitable outcome and maximize detoxification efficiency, cooperators can be periodically reinoculated into the population. Our study investigates the outcome of an evolutionary game combining both public and private goods and demonstrates how environmental parameters can be used to control evolutionary dynamics in practical applications.

## 1 Introduction

Microbes can be detrimental or vital to our health, our environment, and our economy. Much of applied microbiology strives to control these species, by promoting the growth of beneficial species, and suppressing that of harmful ones. We have achieved huge breakthroughs over centuries in eliminating pathogens and preventing dangerous diseases in humans, animals and plants. At the same time, microbes have played important roles in enhancing agriculture and food production (Wolfe and Dutton, 2015), and more recently in the production of biofuels or other chemicals (Antoni et al., 2007; Giri et al., 2020; Quin and Schmidt-Dannert, 2014; Ryan Georgianna and Mayfield, 2012), and in the degradation or “bioremediation” of toxic compounds, such as heavy metals or waste water (Atashgahi et al., 2018; Bertrand et al., 2015; Dixit et al., 2015; Kang et al., 2016; Zaccaria et al., 2020). Despite these exciting advances, these approaches remain only partially successful. In medical microbiology, the emergence of resistant pathogens has led to a life-threatening global health crisis (Tacconelli et al., 2018). On the engineering side, we lack sufficient understanding to maximize the benefits gained from microbes, in terms of production rates or the efficiency of degradation of toxic compounds (Giri et al., 2020).

In all the examples above, there are two obstacles for controlling microbes. First, microbes’ response to changes in their environment, and to the behavior of neighbouring cells may cause the extinction of species that contribute to community function or that protect against pathogens (ecological instability). Second, their large population size and short generation times mean that microbes quickly acquire mutations that can lead to evolutionary instability. Species can thereby gain resistance to antibiotics or lose their ability to perform a desired community function (Akita and Kamo, 2015; Bull and Barrick, 2017; Kumar et al., 1991; Rugbjerg et al., 2018b,a). In the bioremediation of heavy metals, for example, mutants that do not degrade these harmful compounds may invade the original population (Ellis et al., 2007; O’Brien et al., 2014) and exclude it. To optimize the functional efficiency of a microbial system, therefore, we need to consider both ecological and evolutionary dynamics of microbial populations (Schuster et al., 2010).

In this study, we focus in on a bioremediation problem, and develop a mathematical model to investigate the simple case of a single species degrading a toxic compound in a chemostat. We first explore its ecological and evolutionary stability, and second, use this knowledge to optimize the efficiency of toxin degradation over time. In our model, we assume that toxins are harmful to the microbes (e.g., heavy metals), who degrade them by secreting a costly product (e.g., extracellular enzymes). The benefit of degrading toxins is shared by the whole population as, for simplicity, our model does not include spatial structure. Toxin degradation can therefore be regarded as a public goods game (Samuelson, 1954; Broom et al., 2018), as defined in microbiology (Hummert et al., 2014; Smith and Schuster, 2019; West et al., 2007a). It is important to note that not all bioremediation systems correspond to public goods games (e.g., Smith et al. (1998); Röling et al. (2002)), but here we focus on a subset of these systems where the compound to be degraded is toxic to the microbes and its degradation is costly. In such a system, the evolutionary instability of bioremediation is expected (Ellis et al., 2007; O’Brien et al., 2014).

In our model, microbes can adopt one of four strategies: they can be product secretors that pay a cost to contribute to the public good (cooperators) or non-secretors that do not (cheaters) (see O’Brien et al. (2014) for an empirical example). In addition, microbes can be sensitive to the toxins, or can acquire resistance, for example, by activating efflux pumps to expel toxins from within the cell (Blair et al., 2015; Bottery et al., 2016; Rojo-Molinero et al., 2019), or thickening the cell wall. In essence, public good secretion can also be seen as a form of extracellular resistance to the toxin. In other words, here we consider toxin resistance through private or public means, whereby a cell benefits only itself or also the remaining population, respectively.

The population and evolutionary dynamics are then analyzed using evolutionary game theory, where a strategy is considered to be evolutionarily stable if it is not invaded by mutants with another strategy (Maynard Smith and Price, 1973). Evolutionary game theory typically considers the frequencies of strategies (i.e., frequencies sum to one), for example in the replicator dynamics (Cressman and Tao, 2014). Here, however, since toxin concentration decreases with the absolute number or density of degrader microbes (cooperators), our model describes the dynamics of the densities of strategies (as in Hauert et al. (2006, 2008); Gokhale and Hauert (2016)). And since the microbes’ death rate depends on toxin concentration in the environment and their resistance level, our model also includes environmental feedback (as in Gong et al. (2018); Tilman et al. (2020); Weitz et al. (2016)), where each strategy affects the environment differently, and the changing environment affects the fitness of each strategy differently.

We use this model to derive a protocol for optimal toxin degradation. We first show that cooperators that secrete toxin-degrading enzymes are excluded by cheaters that do not, if they have the same level of resistance to the toxin. This recapitulates a well-known result that can be explained by the tragedy of the commons (Hardin, 1968). However, we then show that cooperators can invade a population of cheaters if their level of toxin resistance is different. Since we assume that cheaters are unlikely to acquire double mutations leading to cooperators with a different resistance level, maintaining degradation is only possible if we periodically inoculate these cooperators back into the chemostat. The success of this approach relies on the ability of cooperators to invade cheaters of different resistance. We then calculate the values of the experimentally controllable parameters (inoculation probabilities of cooperators, chemostat dilution rates and inflowing toxin concentrations) that maximize the cumulative efficiency of detoxification.

In sum, our model combines population dynamics, evolutionary dynamics, and environmental feedback to optimize a population-level function. Integrating ecology and evolution into microbial public goods games is increasingly appreciated in microbial applications (Moreno-Fenoll et al., 2017; Sanchez and Gore, 2013). And while optimizing bioremediation is the case study we are considering here, our approach of controlling evolutionary dynamics by changing environmental parameters can be applied to many other microbial functions.

## 2 Model

In our scenario (Fig. 1A), cells can take on one of four strategies depending on whether they produce the enzymes that degrade the toxin (cooperate) or not (cheat), and whether the cells are resistant to the toxin (resistant) or not (sensitive): sensitive cooperator (sCo), sensitive cheater (sCh), resistant cooperator (rCo), and resistant cheater (rCh).

**Figure 1:**
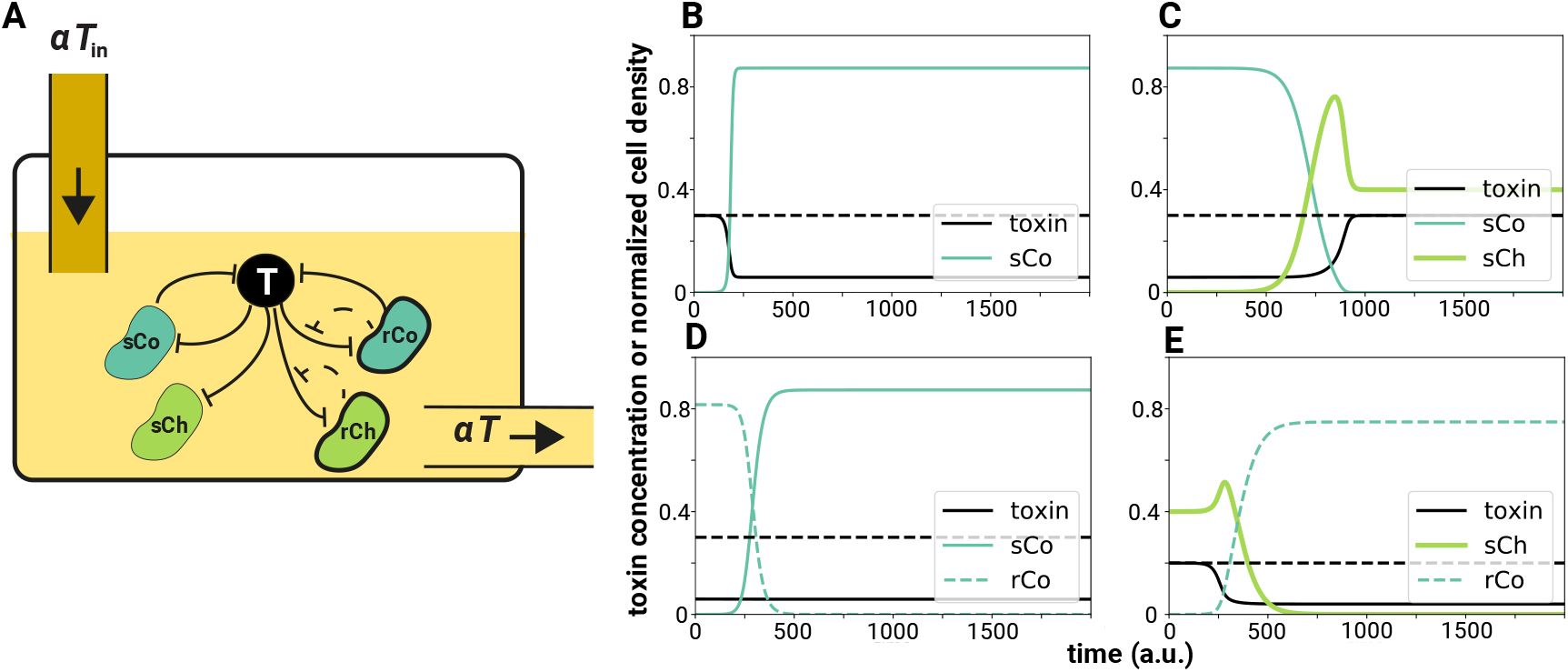
Schematic illustration of the model and examples of the dynamics. (A) In our scenario, a fluid with toxin concentration *T*_in_ flows into the chemostat while the same amount of fluid with toxin concentration *T* flows out. The dilution rate of the chemostat is *α*. Each cell can exhibit one of four strategies: sensitive cooperator (sCo), resistant cooperator (rCo), sensitive cheater (sCh), or resistant cheater (rCh). Cooperators produce enzymes that degrade the toxin, while cheaters do not. The toxin kills cells depending on its concentration, but resistant cells have a lower death rate compared to sensitive cells. Whether the cells are cooperators or cheaters is independent of their resistance level and vice versa. (B-E) Examples of the dynamics in the absence of mutation are shown (a.u. = “arbitrary units”). In each panel, the black solid line represents the toxin concentration *T* and the dashed black line the toxin concentration flowing into the chemostat *T*_in_. Detoxification efficiency at each time-point is proportional to the vertical distance between dashed and solid black lines. Other colored lines represent the cell densities of one of the four strategies (solid dark green: sCo, dashed dark green: rCo, thick lime green: sCh). (B) sCo grows and degrades the toxins. (C) sCo is invaded and excluded by sCh. (D) rCo is invaded and excluded by sCo. (E) sCh is invaded and excluded by rCo. Note that the initial conditions in (C-E) are the stable equilibria of the mono-culture of the resident strategies, while in (B) we begin with a low density of sCo and *T* (0) = *T*_in_. Parameter values are *α* = 0.1, *T*_in_ = 0.2 (E) or 0.3 (otherwise), *f*_max_ = 0.5, *K*_*d*_ = 0.2, *r* = 1, *c*_*d*_ = 0.15, *c*_*r*_ = 0.2 (E) or 0.3 (otherwise), *d*_max_ = 1, *K*_*s*_ = 0.2 (E) or 0.3 (otherwise), *K*_*r*_ = 0.6, and *n* = 1 (E) or 3 (otherwise).

We begin by defining the bacterial population dynamics in our system. The dynamics of the density *x* of each strategy *i* ∈ {sCo, sCh, rCo, rCh} in a chemostat is defined by growth, death and dilution out of the chemostat:

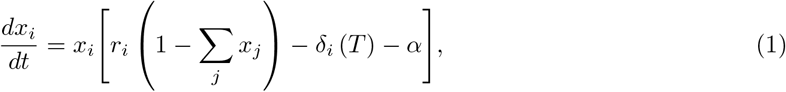

where *r*_*i*_ is the intrinsic growth rate of strategy *i* due to nutrients that are not explicitly defined in the model, *δ*_*i*_ (*T*) is the death rate of strategy *i* given toxin concentration *T*, and *α* is the dilution rate. The total densities of the four strategies ∑_*i*_ *x*_*i*_ should be lower or equal to one in Eq (1); i.e., the carrying capacity is equal to one in the absence of death or dilution. In this formulation, a useful proxy for fitness is the ratio between intrinsic growth and death at a given toxin concentration, whether death is by toxin or by dilution:

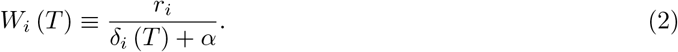

At an equilibrium, *W*_*i*_(*T*) = 1/ (1 − ∑_*j*_*x*_*j*_) should be satisfied for any strategy *i* that exists in the chemostat (*x*_*i*_ > 0). In addition, when *W*_*i*_(*T*) > *W*_*j*_(*T*), strategy *i* increases faster or decreases slower than strategy *j*. For simplicity, this basic model assumes that strategies cannot mutate into each other. We extend it to include mutations in Appendix 5.

First, the intrinsic growth rates in this model differ depending on the costs each strategy pays. Cooperators pay a cost, *c*_*d*_, for producing degrading enzymes, which are regarded as a public good since they reduce environmental toxicity and the death rate of all cells independently of their strategy. In addition, toxin resistance carries a cost, *c*_*r*_. Such fitness costs, where resistant cells have lower fitness than sensitive ones in the absence of toxins, have been observed in many species (Andersson and Levin, 1999; Andersson and Hughes, 2010; San Millan and MacLean, 2017). In contrast to the production of degrading enzymes, however, where all cells benefit from decreased toxicity, the evolution of resistance can be regarded as an investment into a private good, where only the resistant cells themselves benefit. Assuming that the costs are additive, the intrinsic growth rate *r*_*i*_ of each strategy is defined as follows:

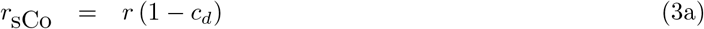

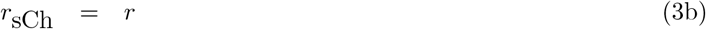

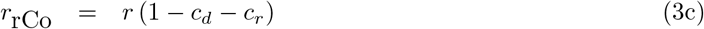

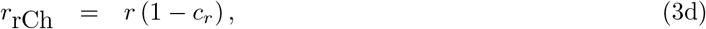

where *r* is the maximum intrinsic growth rate.

Cellular death rate *δ*_*i*_(*T*) increases with toxin concentration *T*, and is represented by a Hill equation as is common in models of death by drugs (Chou, 2006):

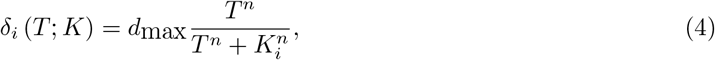

where *d*_max_ is the maximum death rate, *K*_*i*_ is the half maximal toxin concentration of strategy *i*, and *n* is the Hill coefficient, which determines the steepness of the function. Resistance can be modeled either by increasing *K*_*i*_ or decreasing *n* (Sampah et al., 2011). Here we assume that resistant cells have a larger *K* than sensitive cells: *K*_*r*_ > *K*_*s*_, such that they reach *d*_max_ at a higher toxin concentration than the sensitive cells. Note that the toxin concentration *T* changes over time, as described below.

Due to the dilution in the chemostat, a proportion of cells of each strategy *i* flows out of the chemostat. The dilution rate into and out of the system is denoted by *α*.

To describe the population dynamics of each strategy, however, it is necessary to also formulate the dynamics of the toxin concentration because it affects the microbes’ death rate, and because the toxin concentration changes over time as cooperators detoxify it. The dynamics of the toxin concentration *T* in the chemostat are defined by the concentration flowing into and out of the chemostat, and detoxification by cooperators:

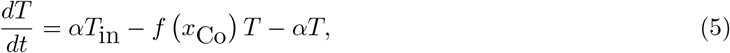

where *T*_in_ is the toxin concentration flowing into the chemostat, and *f*(*x*_Co_) is the degradation rate which is assumed to follow a Michaelis-Menten function:

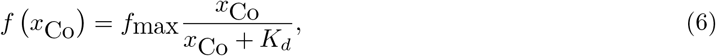

where *x*_Co_ = *x*_sCo_ + *x*_rCo_; i.e., the sum of sensitive and resistant cooperators. Whether cooperators are resistant or sensitive has no impact on toxin degradation. *K*_*d*_ represents the density of cooperators *x*_Co_ that gives half the maximum of *f*(*x*_Co_). All parameters of the model are listed in Table 1.

**Table 1:** List of parameters

| Notation | Range | Description |
| --- | --- | --- |
| $\alpha$ | $(0, 1]$ | dilution rate of the chemostat |
| $r$ | $(0, 1]$ | maximum intrinsic growth rate of the microbe |
| $c_d$ | $(0, 1]$ | cost of cooperation (production of the degrading enzyme) |
| $c_r$ | $(0, 1]$ | cost of resistance to the toxin |
| $d_{\max}$ | $(0, 1]$ | maximum death rate by toxin |
| $K_s$ | $[0, K_r)$ | half maximal effective toxin concentration of the sensitive cells |
| $K_r$ | $(K_s, 1]$ | half maximal effective toxin concentration of the resistant cells |
| $n$ | $(0, \infty]$ | Hill coefficient |
| $T_{\text{in}}$ | $(0, 1]$ | toxin concentration flowing into the system |
| $f_{\max}$ | $(0, 1]$ | maximum degradation rate of the toxin |
| $K_d$ | $(0, 1]$ | half maximal effective cooperator density of the degradation rate |
| $\mu_1$ | $[0, 1]$ | mutation probability in the function of detoxification |
| $\mu_2$ | $[0, 1]$ | mutation probability in the resistance level |
| $m_1$ | $[0, 1]$ | inoculation probability of sCo |
| $m_2$ | $[0, 1]$ | inoculation probability of rCo |

As the goal of this study is to maximize the efficiency of detoxification, we define detoxification efficiency *ϕ* as the difference between the toxin concentration flowing into and out of the chemostat multiplied by the dilution rate:

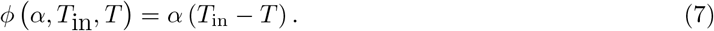

With this definition, *ϕ* is proportional to the amount of detoxified liquid and is composed of the degree of detoxification and the amount of liquid flowing out of the chemostat. Although this equation gives the efficiency at any time *t*, we mainly focus on the efficiency at an equilibrium.

## 3 Results

### 3.1 All strategies can persist in mono-cultures with no mutation

We first analyse whether cooperators and cheaters can persist in mono-culture. Remember that only cooperators produce public goods that degrade the toxin and thereby increase the survival of all cells in the chemostat. When a few cooperators (either sensitive or resistant, 0 *< *x*_i_*(0) ≪ 1, *i* ∈ {sCo, rCo}) are introduced into the system, they increase and converge to an equilibrium of positive density (Fig. 1B) if and only if

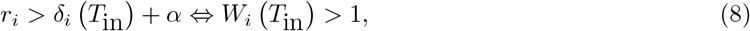

where *i* is the focal strategy (see Appendix 1 for derivation). By solving *dT/dt* = 0 and *dx*_*i*_*/dt* = 0, one can find a trivial equilibrium (*x*_*i*_ = 0) and one or more non-trivial equilibria 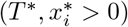 that should satisfy:

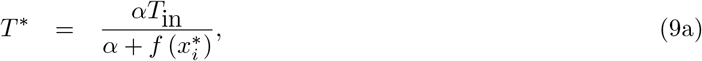

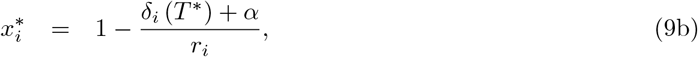

which we can calculate numerically using Newton’s method.

In the absence of cooperators, we assume that the toxin concentration is equal to the incoming toxin because cooperators are the only degrader cells. In the case of a mono-culture of cheaters then, *T*\* = *T*_in_ regardless of cell density, and their density at a stable equilibrium 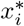, *i* ∈ {sCh, rCh} is positive and given by Eq (9b) if inequality (8) holds (see Appendix 1 for details). From now on, we focus on conditions where cell density converges to a positive value in mono-culture (i.e. where inequality (8) holds).

### 3.2 Cheaters can invade a population of cooperators

Next, we ask what happens when a cheater mutant invades a population of cooperators at its non-trivial equilibrium or vice versa. Cheater mutants can invade and exclude a population of cooperators at any toxin concentration and independently of whether they are both sensitive or both resistant, as long as the resident and mutant have the same resistance levels (e.g., sCo and sCh) (Figs. 1C, and 2A). In contrast, cooperators are unable to invade a population of cheaters of the same resistance level at any toxin concentration. These findings recapitulate the classical result that cheaters will always dominate in a well-mixed environment (West et al., 2007b) because cooperators pay a cost for producing degrading enzymes, but are as sensitive to the toxin as cheaters. In other words, the tragedy of commons (Hardin, 1968) occurs in this case.

**Figure 2:**
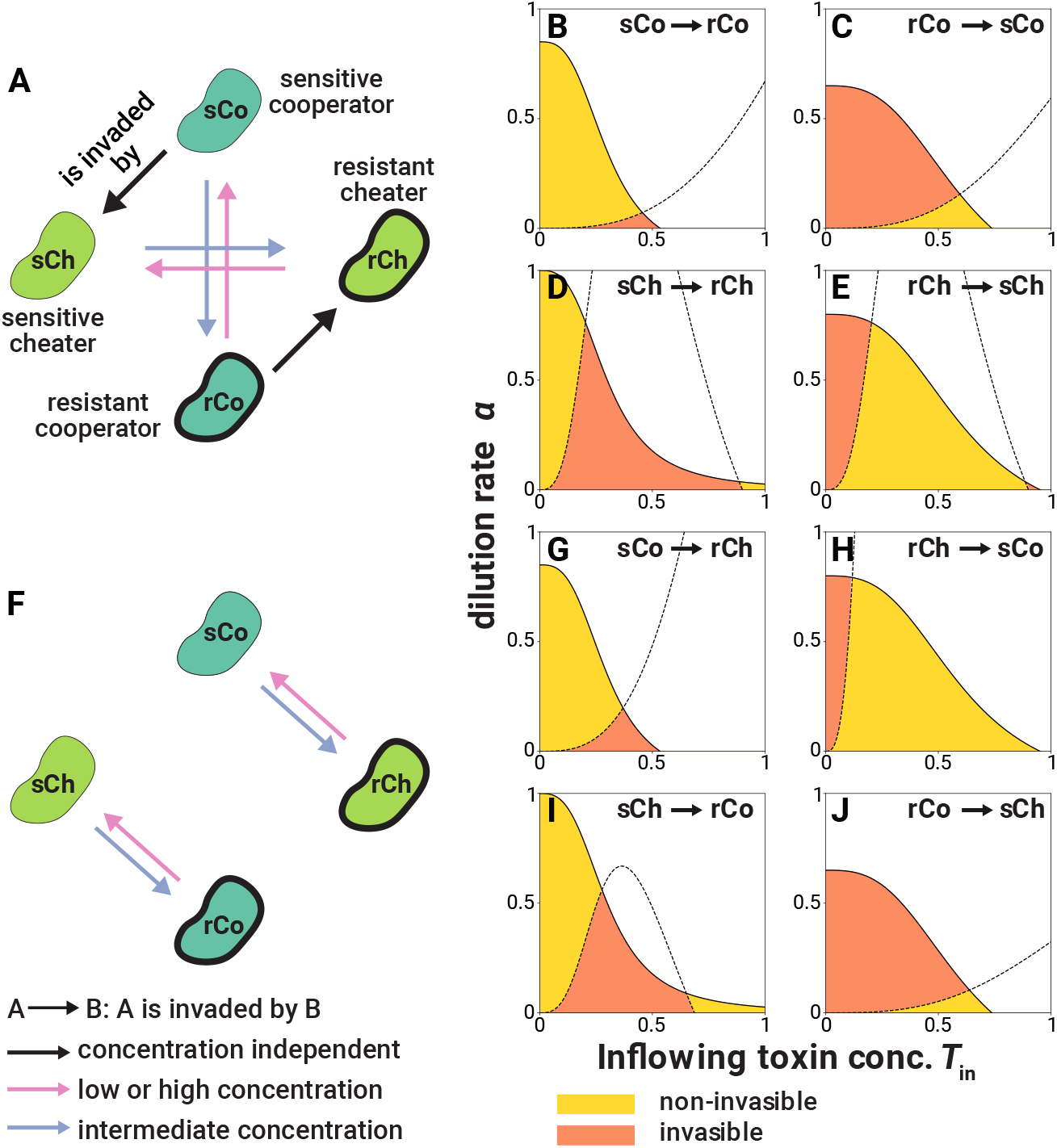
Short-term evolutionary dynamics of pairwise invasions. (A, F) Diagrams of pairwise invasion analysis when a single-mutation (A) or double-mutations (F) occur. A → B represents that A is invaded by B, and the color of each arrow shows the condition for successful invasion. Black arrows represent successful invasion regardless of the toxin concentration, while pink and blue arrows represent toxin-concentration-dependent invasion (that invasion succeeds when the toxin concentration is low or high, or when the toxin concentrations are intermediate, respectively). (B-E, G-J) Examples of an invasion state space for each pair of strategies. A → B represents that A is a resident strategy and B is an invader. Pairs of sCo and sCh, and rCo and rCh are omitted since cheaters are fitter than cooperators in these pairs, regardless of the parameter values. In the yellow areas, the resident strategies are not invaded by the invaders, while the invasion succeeds in the orange areas. Each solid line is a boundary under which the resident strategy persists in mono-culture. The dashed line in each panel represents where the fitness proxy *W* of resident and invader strategies are equal at an equilibrium reached in a mono-culture of the resident strategy. Note that residents can coexist with invaders under certain conditions (see Appendix 3). Parameter values in (B-E, G-J) are *f*_max_ = 0.5, *K*_*d*_ = 0.2, *r* = 1, *c*_*d*_ = 0.15, *c*_*r*_ = 0.2, *d*_max_ = 1, *K*_*s*_ = 0.3, *K*_*r*_ = 0.6, and *n* = 3. Note that panels (B-E, G-J) are examples of the state space given the parameter values; different parameter values will show different invasion landscapes.

### 3.3 Toxin concentration determines invasion of sensitive and resistant cells

For mutants that differ in their private resistance level (e.g., sCo and rCo), the toxin concentration determines whether invasion succeeds or not (Figs. 2B-E). Intuitively, this is because the benefit of being resistant to the toxin is quite low when its concentration is very low. Similarly, when toxin concentration is very high, the death rate of the resistant strain is close to that of the sensitive strain and too high to compensate for the cost of private resistance. Under these conditions, sensitive cells can invade a population of resistant ones (Fig. 1D). Instead, resistant cells can invade a population of sensitive cells when the toxin concentration in the chemostat is intermediate. Two strategies that differ only in their resistance level never stably coexist (see Appendix 3 for derivation).

### 3.4 Cooperators can invade and coexist with cheaters of different resistance levels

Thus far, we have considered whether mutants can invade a resident population that differs in only one trait, their private or their public resistance (i.e. cooperative toxin degradation). While we assume that the time to reach an equilibrium following a single mutant invasion is shorter than the time for a second mutation to occur, we nevertheless explore invasions by such double mutants here (Fig. 2F). Depending on the concentration of toxins, rCo and sCh can invade each other’s populations, as can sCo and rCh (Figs. 2G-J). This dependency on toxicity follows the same logic as for the invasion of a resistant mutant into a sensitive population described above: when the toxin concentration is intermediate, rCo have a much lower death rate than sCh, and the benefit of resistance exceeds the sum of the cost of cooperation *c*_*d*_ and resistance *c*_*r*_. If, on the other hand, the toxin concentration is either too low or too high, resistance to the toxin does not provide enough of an advantage to overcome its cost, leading instead to the invasion of sCh into a population of rCo. The same logic, albeit with different thresholds can explain the invasion of sCo into rCh and vice versa (Figs. 2G-J). In sum, invasions of double mutants into resident populations that differ in both public and private resistance depends on toxin concentrations (see Appendix 2).

Once a mutant has invaded, whether it will coexist with the resident population is unclear because, as cooperators increase, toxin concentration decreases, which changes the fitness landscape. In other words, increasing cooperator density can decrease the fitness difference between cooperators and cheaters. In Appendix 3 we show that cooperators and cheaters of different resistance levels (e.g. rCo and sCh) can indeed stably coexist at certain parameter ranges. Nevertheless, these two coexisting strains can then be invaded by cheaters with different resistance (e.g. rCh), which excludes the other two strategies (see Appendix 4). In other words, cooperators are never evolutionarily stable because they can be invaded and excluded by cheaters of the same resistance level, as we show next.

### 3.5 In the long-term, cooperators are unlikely to be maintained

Having analyzed the outcomes of the invasion of all mutants in the short-term (Figs. 2A and F), we can now predict how the population in the chemostat will change in the long-term. Regardless of which type we start with, as cells mutate, the population will transition between different genotypic “states”, which can be represented by the state transition diagram in Fig. 3. The probability of cooperators mutating into cheaters or vice versa is given by *μ*_1_, while *μ*_2_ is the probability to change the level of resistance. For simplicity, we assume that these mutation probabilities are very small. Once a mutation occurs, we use Eq (1) as outlined above to take us to the following equilibrium state. We assume that no further mutations will occur before the equilibrium is reached, but relaxing this assumption does not alter the overall dynamics (Appendix 7). For example, sCh will appear in the population of sCo with probability *μ*_1_ (1 − *μ*_2_) and exclude it. Then, rCh can appear in the population of sCh with probability (1 − *μ*_1_) *μ*_2_, but may invade or not, depending on the toxin concentration (Fig. 3).

**Figure 3:**
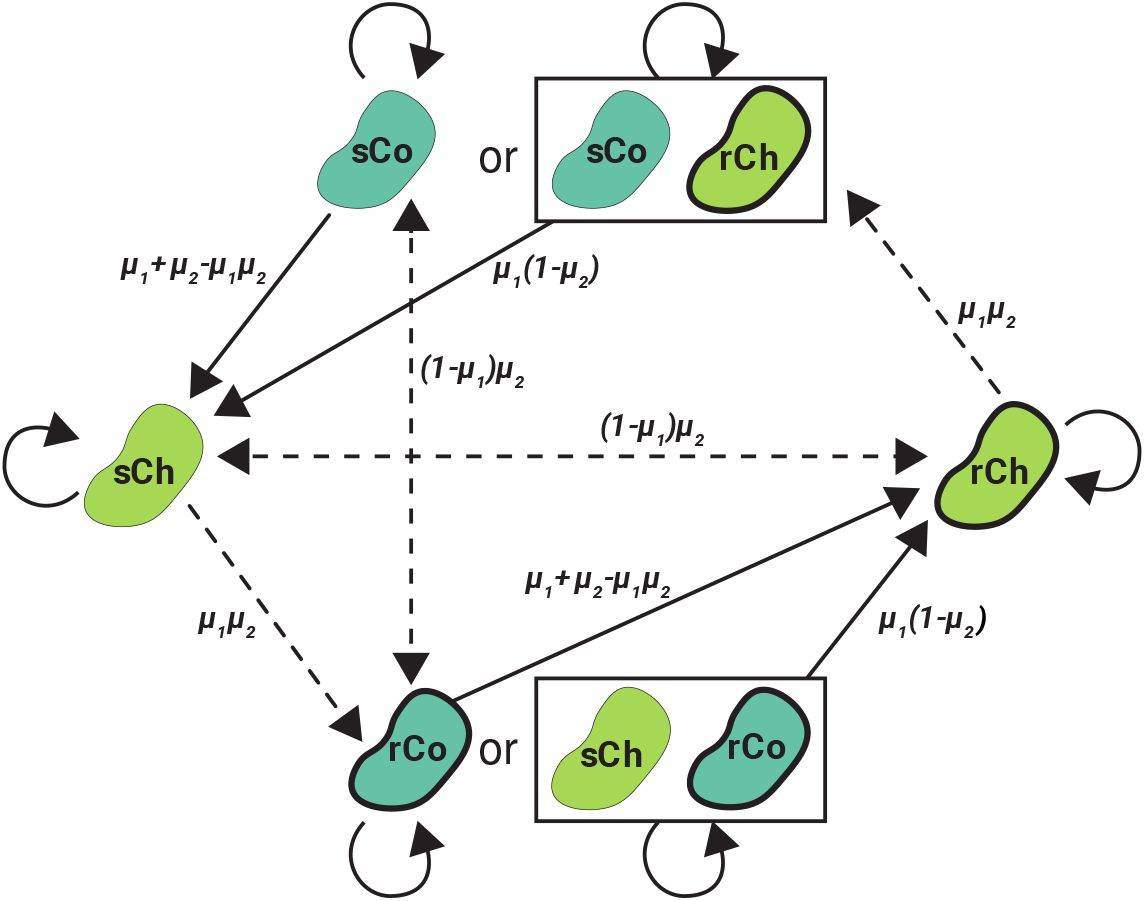
Schematic illustration of the state transitions for long-term evolutionary dynamics. Arrows represent state transitions resulting from natural selection. Solid arrows show transitions that are independent of toxin concentration, and dashed arrows transitions that depend on toxin concentration, here depicted for a given *T*_in_ and *α*. The transition from sCo to rCo and vice versa do not occur if cooperators coexist with cheaters. Values along the arrows represent state transition probabilities.

Although in principle cooperators can invade a population of cheaters that differ in the level of resistance (e.g., rCo can invade sCh), (i) invasion success depends on *α* and *T*_in_, (ii) double mutations are expected to be rare (*μ*_1_*μ*_2_ is close to 0) and (iii) if a cheater mutant of the same resistance as the cooperator invades (e.g. rCh), it will dominate the population and replace the cooperators. Accordingly, it is very difficult to maintain cooperators in the chemostat due to natural selection. This brings us to one of the main findings of the study: even though cooperators and cheaters can coexist under some conditions, to maintain costly microbial detoxification, it is necessary to inoculate cooperators manually and to change *α* and *T*_in_ to favor their survival. Crucially, though, because cooperators are able to invade cheaters of opposite sensitivity, these inoculations can maintain cooperators – and thereby detoxification – in the short-term. In the following sections, we show how to control the values of *α* and *T*_in_ and inoculation probabilities to maximize the efficiency of detoxification.

### 3.6 Culture conditions can be controlled to optimize detoxification efficiency

Ultimately, our goal is to maximize the efficiency of detoxification *ϕ*, which depends on the absolute abundance of the two types of cooperators in the chemostat. In turn, these abundances can be controlled by changing the culture conditions through two parameters: the dilution rate *α* and the toxin concentration flowing into the chemostat *T*_in_.

To maximize the objective function in Eq (7), we consider three stable equilibrium states with different toxin concentrations flowing out of the chemostat: (i) *T* = *T*_in_ when only cheaters are present, regardless of the values of *α* and *T*_in_, (ii) 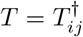 when cooperators *i* coexist with cheaters *j*, which have different resistant levels, and (iii) 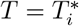 when only one type of cooperators *i* is present. In the latter two cases, one can calculate the equilibria (analytically or numerically) and their corresponding detoxification efficiency *ϕ* for each culture condition (values of *α* and *T*_in_). We can then find the optimal culture conditions that maximize this efficiency (Fig. 4), although the equilibrium can be ecologically unstable for some parameter values.

**Figure 4:**
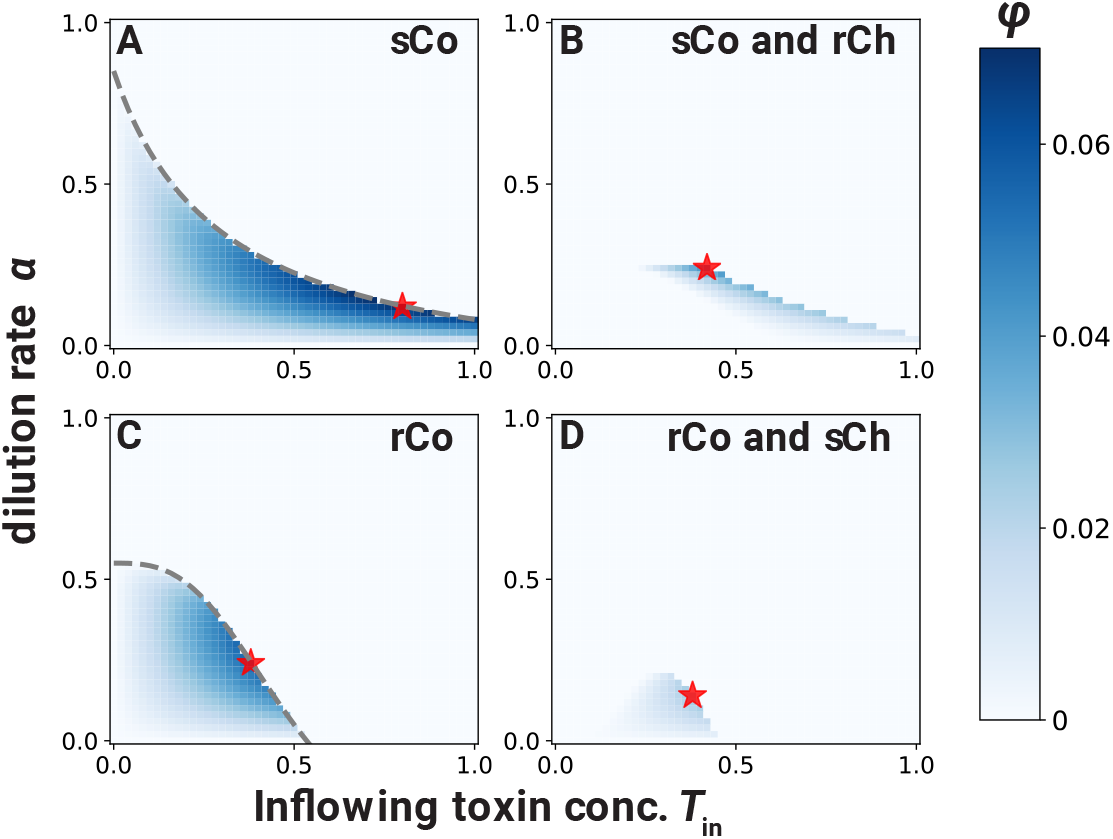
Optimal detoxification at each equilibrium. The efficiency of detoxification at an equilibrium state *ϕ(α, T*_in_*, T*) given *α* and *T*_in_ is represented by color in each panel: (A) only sCo, (B) coexistence of sCo with rCh, (C) only rCo, and (D) coexistence of rCo with sCh. The red stars represent the maximum efficiency of detoxification in each. In the areas above the dashed gray lines in each two left panels, the cooperator cannot persist in mono-culture (inequality 8). Parameter values are *f*_max_ = 0.5, *K*_*d*_ = 0.2, *r* = 1, *c*_*d*_ = 0.15, *c*_*r*_ = 0.3, *d*_max_ = 1, *K*_*s*_ = 0.3, *K*_*r*_ = 0.5, and *n* = 1 (A-B) or *n* = 3 (C-D). We used different values of *n* in the top and bottom rows to allow cooperators to coexist with cheaters with a different level of resistance.

Intuitively, the maximum efficiency is larger in a mono-culture of cooperators than in a co-culture of cooperators and cheaters of different levels of resistance (see Appendix 6). If cheaters can be excluded from the population by changing *α* and *T*_in_, the optimal strategy for cultivation is (i) to exclude the cheaters by adjusting the culture conditions, and then (ii) to change the culture conditions to maximize the productivity of a mono-culture of cooperators.

### 3.7 Inoculating cooperators to optimize detoxification efficiency

Above, we showed that even though they are unlikely to appear by double mutation, cooperators can invade a population of cheaters if their level of resistance is different (Fig. 3). Instead of waiting for these mutants to arise naturally, it would be more efficient to manually inoculate cooperators into the population, and to change *α* and *T*_in_ to allow them to invade successfully and to exclude the cheaters. Assuming that we cannot observe the prevalence of each strategy at will, the problem is how often to inoculate sensitive or resistant cooperators to maximize detoxification efficiency over time. If cooperator inoculation probabilities are too small, cheaters will dominate the population, leading to a detoxification efficiency of zero. If they are too large, we may inoculate cooperators unnecessarily (e.g., sCo into a monoculture of sCo) or when they cannot invade (e.g., sCo into a mono-culture of sCh). Such unfavorable inoculations can be costly because they can require a higher in-flowing toxin concentration *T*_in_, and result in reduced detoxification efficiency for some time.

To optimize cooperator inoculation probabilities *m*_1_ and *m*_2_ for the sensitive or resistant cooperators, respectively, we consider population state transitions as a Markov chain with discrete time-steps *s*, and find the values of *m*_1_ and *m*_2_ that maximize the total amount of toxin degradation (Figs. 5A, and A.9). Transitions in this Markov chain model can occur either due to mutations at probabilities ***μ*** = (*μ*_1_*,μ*2) or inoculations at probabilities ***m*** = (*m*_1_,*m*_2_), resulting in the transition matrix *P* (***m***; ***μ***). The state distribution vector ***π***(*s*) = (*π*_*i*_(*s*)) where *π*_*i*_(*s*) is the probability that the population is in state *i* at time step *s* (∑_*i*_ π_*i*_(*s*) = 1 for *s* = 0, 1, …, ∞). Because the Markov chain is ergodic when cooperators can exclude cheaters that differ in their resistance level (Fig. 5A, but see Appendix 7 for a case where cooperators cannot exclude cheaters), the probability distribution of the population states converges to a unique stationary distribution ***π***^*^ in the limit of *s* → ∞, regardless of the initial distribution ***π***(0):

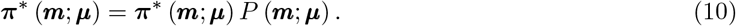

**Figure 5:**
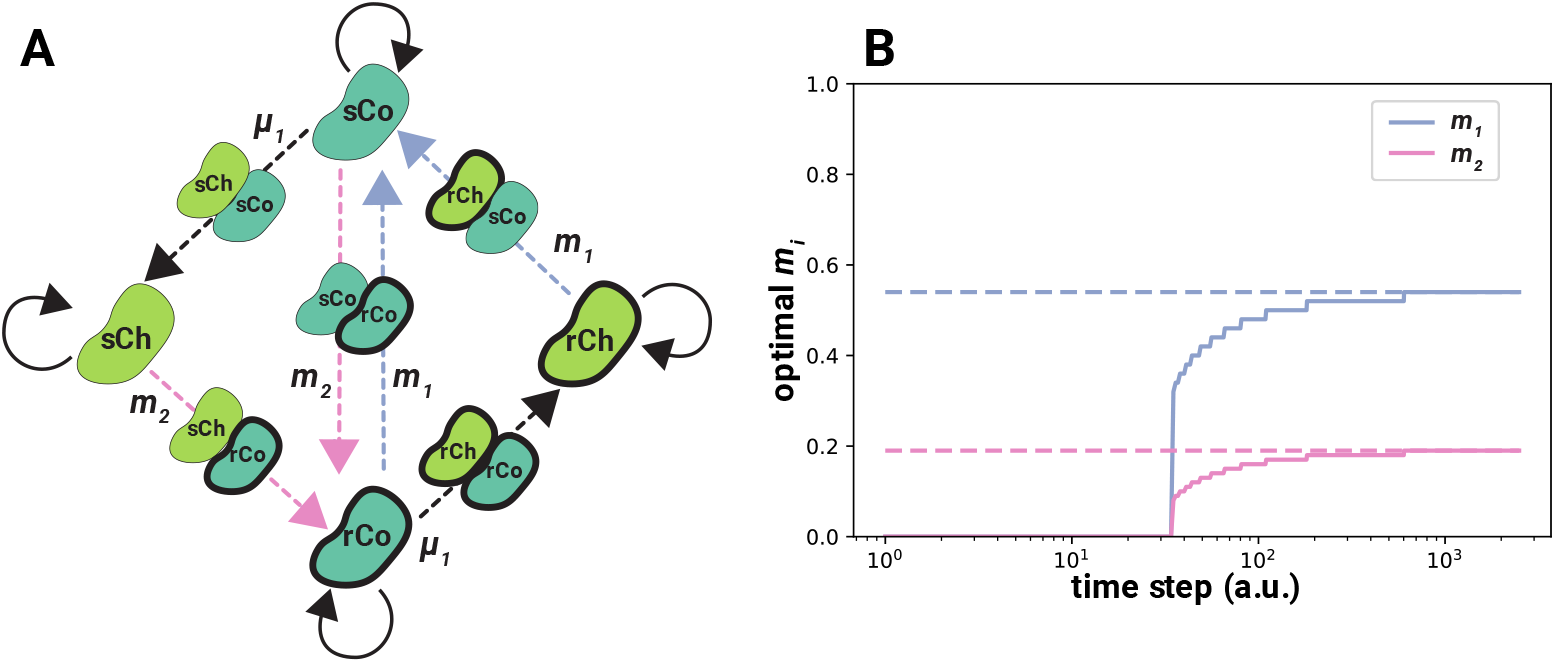
Optimization problem assuming that resistance mutations do not occur. (A) A simplified schematic illustration of the state transition diagram when *μ*_2_ = 0 and cooperators can exclude cheaters of different resistance levels. A *→* B represents a transition from state A to state B. Colored arrows represent transitions that occur through the introduction of sensitive and resistant cooperators in blue and pink, respectively. Dashed arrows indicate transient states where two strategies coexist. The full diagram containing 14 states is shown in Fig. A.9. (B) The optimal values of *m*_1_ and *m*_2_ (the blue and pink solid lines, respectively) which maximize the cumulative efficiency defined by Eq (A.67) calculated using Dynamic Programming. The two dashed lines represent the values of *m*_1_ and *m*_2_ which maximize Eq (11). The initial state of the population is a mono-culture of sCo. Around 1,000 time steps, the optimal values of *m*_1_ and *m*_2_ for Eq (A.67) converge to the values which maximize Eq (11). In practice, the detoxification efficiency at each of the 14 states of the model as well the mutation probabilities would be experimentally measured and plugged into the model to calculate the values of *m*_1_ and *m*_2_ that would maximize cumulative efficiency. The plot above was generated using the following fictitious values, as an illustration: *μ*_1_ = 0.01, *μ*_2_ = 0, ***ϕ*** = {0.4, 0.2, 0, 0.15, 0.3, 0.15, 0, 0.2, 0.35, 0.35, 0.3, 0, 0.2, 0}. See Appendix 7 for more detail.

Even though the transitions are probabilistic, we assume that the establishment of strategies following mutation or inoculation (i.e., short-term dynamics) is deterministic. Relaxing this assumption by introducing an establishment probability *E* into the transition matrix *P* (***m***; ***μ***) does not change the ergodicity of the Markov chain, and we arrive at the stationary distribution in the same manner. In Appendix 7 we show how to calculate the expected cumulative efficiency of detoxification Φ from the beginning of the cultivation to time step *s* (defined in Eq (A.67)). This calculation is somewhat cumbersome, but for large *s*, Φ is approximately proportional (see Appendix 7) to the expected efficiency of detoxification at the stationary distribution ***π***^*^:

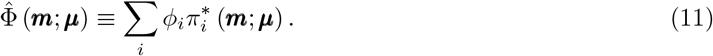

To maximize the expected cumulative detoxification efficiency Φ, therefore, we can calculate ***m*** that maximizes Eq (11). In Box 1 we show an example of how to use this approach in practice.

##### Box 1: An example of optimizing detoxification efficiency

Imagine that we have set up the experimental system described, and would like to compute the optimal inoculation probabilities. We first need to define the Markov chain to make predictions, and second, we need to experimentally measure the parameters of our bacterial strains, in particular their degradation efficiency *ϕ*_*i*_ at each of the different states *i* of the Markov chain.

To establish the Markov chain, we begin with a few simplifying assumptions: (i) that *μ*_2_ = 0, such that cooperators can only invade a population of cheaters that differ in the level of resistance by inoculation, and (ii) that mutations and the manual inoculation of a cooperator strategy can occur only in a mono-culture (i.e., at most two strategies can exist simultaneously in the population). We further assume that the parameters are in a range where at certain (*α, T*_in_), (iii) sCo and rCo can mutually exclude each other, and (iv) sCo and rCo can exclude rCh and sCh, respectively. Under these assumptions (we relax (i) below and (ii)-(iv) in Appendix 7), the Markov chain consists of at least 14 states: four mono-culture situations, six transient situations where two strategies coexist, and four situations where the introduction of cooperators is unfavorable (see Fig. A.9 for the diagram). A simplified schematic of this model is shown in Fig. 5A.

By experimentally measuring mutation probabilities ***μ*** and detoxification efficiencies ***ϕ*** = (*Cϕ*_*i*_) for each state *i*, where *C* is a positive constant to change the time scale of *ϕ*_*i*_ into a discrete time step, we can calculate the probability distribution ***π*** (*s, **m***) = (*π*_*i*_ (*s*, ***m***)) as a function of time step *s* and inoculation probabilities ***m*** = (*m*_1_*, m*_2_). Then, Dynamic Programming (DP) provides the optimal ***m*** that will maximize the cumulative expected detoxification efficiency Φ when the experiment finishes at time step *s*. Starting from a mono-culture of sCo, the two solid lines in Fig. 5B represent *m*_1_ and *m*_2_ provided by DP given some fictitious yet reasonable values of ***μ*** and ***ϕ***. At first, the optimal values of *m*_1_ and *m*_2_ are zero, because the state of the population is most likely to be a mono-culture of sCo, in which case inoculating cooperators would be pointless. However, as time passes, mutations will arise, and the population is likely to transition to a state of sCh mono-culture; then, the optimal values of *m*_1_ and *m*_2_ increase. At about 1,000 time steps, the optimal values of *m*_1_ and *m*_2_ converge to the values which maximize detoxification efficiency at the stationary distribution 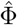 described by Eq (11). When *μ*_2_ > 0 (relaxing assumption (i)), the number of states increases and the state transition diagram becomes more complex. As long as the Markov chains is ergodic, however, it is possible to find the stationary distribution ***π***^*^ and the optimal values of *m*_1_ and *m*_2_ that maximize Eq (11). We show how to find the optima for non-ergodic Markov chains in Appendix 7.

## 4 Discussion

In this study, we have shown how to control the ecological and evolutionary dynamics of a microbial population growing in a chemostat in order to optimize the bioremediation of a toxic liquid. Public goods games where cooperators increase the growth rates of surrounding cells at a cost to themselves have been extensively studied, both empirically and theoretically (Allen et al., 2016; Hauert et al., 2006, 2008; Griffin et al., 2004; Gokhale and Hauert, 2016; Sanchez and Gore, 2013). Rather than increasing the growth rate of others, cooperators in our model degrade toxic compounds, which decreases the death rate of surrounding cells (O’Brien et al., 2014). This scenario enables us to introduce the evolution of resistance to the toxin, e.g. through efflux pumps, as a private good, which we base on studies of drug-dose effect and resistance to it (Chou, 2006; Sampah et al., 2011). Unsurprisingly, cheaters always exclude cooperators with the same private resistance level because detoxification is costly (West et al., 2007b). We show, however, that because the benefit of private resistance depends on the toxin concentration in the chemostat, cooperators can invade a population of cheaters that differ in their private resistance. The co-occurrence of two strains that differ both in their degradation ability as well as their resistance level is unlikely to suddenly arise by mutation, especially if we assume that mutations are rare. To maintain the degradation of toxins, therefore, it is necessary to periodically inoculate cooperators into the chemostat while changing the dilution rate and in-flowing toxin concentration to guarantee invasion success.

Optimal values for these parameters (cooperator inoculation probabilities, dilution rate, and in-flowing toxin concentration) that maximize the detoxification efficiency of the system can be calculated using our model. As input, the model requires experimental measurements of growth and death rates of the chosen microbe and its mutants (i.e., intrinsic growth rate of each strategy *r*_*i*_, maximum death rate *d*_max_, Hill coefficient *n*, median-effect toxin concentrations *K*_*s*_, *K*_*r*_, and degradation efficiencies *ϕ* of cooperators).

Our model and its results can also apply to problems other than bioremediation that involve survival in toxic environments. In essence, we are studying the evolutionary dynamics of public resistance (which is cooperative) and private resistance (which is not). Consider, analogously, two types of antibiotic resistance mechanisms: public mechanisms are costly and benefit the producing cell as well as its neighbors such as extracellular secretion of antibiotic-degrading enzymes (e.g. β-lactamases (Yurtsev et al., 2013)), and private resistance mechanisms only benefit the producing cells, such as efflux pumps. The evolutionary dynamics in a scenario whereby cells can switch between these different resistance mechanisms and being sensitive to the antibiotic correspond to Figs. 2A and F. In this case, however, an objective function would aim to minimize rather than maximize the densities of the most resistant strains. Another interesting aspect is that the benefits of resistance depend on toxin concentration in the chemostat, which is affected by the density of cooperators and the toxin concentration flowing into the chemostat. In other words, the public goods game affects the benefit of the private goods. This is why cooperators can invade a population of cheaters when they differ in their resistance level (Fig. 2F).

Of course, our model relies on a number of assumptions and focuses only on a subset of possible bioremediation systems. First, we only consider extracellular toxin degradation (e.g. by enzyme secretion), while toxins can also be degraded inside cells (O’Brien and Buckling, 2015). For intra-cellular degradation, a different functional form of detoxification *f* (*x*_Co_) would be necessary, but we expect similar results as long as this function increases monotonically with the density of cooperators. Indeed, the invasion analysis is independent of the form of *f*(*x*_Co_). Similarly, we assume that toxins kill the microbes and that their degradation does not contribute to growth. In reality, many compounds that are undesirable for humans are instead used as substrates by microbes (Atashgahi et al., 2018). This latter case is simpler than the one we consider here, since detoxification is no longer cooperative and there is no risk of cheaters arising and collapsing the system. Finally, detoxification may carry a negligible cost, for example if it the toxic compound is neutralized by a change in pH, which occurs naturally due to a microbe’s metabolism.

Another issue is how to define detoxification efficiency *ϕ*. Rather than Eq (7) one could, for example, define *ϕ* as the time needed for the toxins to decrease to a negligible concentration. This would change the optimal culture conditions *α* and *T*_in_, but not the procedure to find the optimal introduction probabities of cooperators ***m***, which are independent of the formulation of *ϕ*. Our model also fixes some parameters, such as the Hill coefficient *n*, which can evolve in reality (Sampah et al., 2011). Similarly, the cost of resistance *c*_*r*_ can decrease over time due to compensatory evolution (Andersson and Hughes, 2010; San Millan et al., 2014). Allowing these parameters to evolve would make it more difficult for sCo to invade rCh because the relative fitness of rCh will increase.

We also assume that our system is well-mixed and that there are no spatial gradients within the chemostat. Spatial structure, for example, whereby detoxifying enzymes diffuse slowly through the chemostat and have a patchy distribution can favor the coexistence of cooperators and cheaters (Allison, 2005). Indeed, previous empirical bioremediation studies have reported coexistence of cooperators with cheaters (Ellis et al., 2007; O’Brien et al., 2014). Theoretically, this may be due to a difference of resistance levels between cooperators and cheaters as we show here, but a simpler explanation would be the presence of spatial gradients. Relaxing the assumption of a perfectly well-mixed chemostat would make thepersistence of cooperators easier. It may also increase the public benefit of toxin resistance, which we have considered to be private here (Rojo-Molinero et al., 2019).

Finally, there may be other ways of periodically introducing cooperators. Experimentally, our constant inoculation probabilities represent a situation where stock strains of cooperators would be manually added into the chemostat. If instead, multiple chemostats are running in parallel, another way of introducing cooperators would be to exchange certain amounts of fluids between chemostats. Ecologically, this would correspond to migration among patches, and the optimal migration probabilities would depend on the probability distribution of the different strategies in each chemostat. Comparing the optimal introduction probabilities and the cumulative efficiency of detoxification between the model presented here and a multi-chemostat system is left for future work.

In summary, we have combined an ecological model with evolutionary game theory to develop a protocol for the control and optimization of a bioremediation system by microbes, and guard it against collapse through the emergence of cheaters. More broadly speaking, our scenario motivates the integration of important elements from ecological models, such as population densities and environmental feedback, into evolutionary game theory. In essence, our model can be adapted to any practical applications involving costly microbial traits, where manipulating environmental conditions can be used to control evolutionary dynamics. Achieving this will allow us to better anticipate evolutionary change in microbial systems that we strive to control, whether this involves increasing toxin degradation as we have shown here, the production of public goods such as useful chemicals, or eliminating antibiotic resistant pathogens.

## 5 Data availability

Python code and csv files used for simulations and analytic calculations are available at GitHub, but we cannot show the url for double-blinding.

## Acknowledgements

We thank Masato Yamamichi and Xiang-Yi Li for useful feedback on the earlier versions of the manuscript. SS is funded by the Nakajima foundation and the University of Lausanne and SM by European Research Council Starting Grant 715097.

