## Supplementary information for "Controlling evolutionary dynamics to optimize microbial bioremediation"

### Appendix 1 Mono-culture analysis

In this section, we analyze the mono-culture of each of the four strategies. From Eqs (1) and (5) in the main text, the null-clines for strategy $i$ are written as follows:

$$T = \frac{\alpha T_{\text{in}}}{\alpha + f\left(x_{\text{Co}}\right)} \tag{A.1a}$$

$$x_i = 1 - \frac{\delta_i\left(T\right) + \alpha}{r_i} \tag{A.1b}$$

$$x_i = 0. \tag{A.1c}$$

Eq (A.1a) represents a decreasing function of $x_{\text{Co}}$ (the density of the cooperators; i.e., in the mono-culture $x_{\text{Co}} = x_i$ where $i \in \{\text{sCo}, \text{rCo}\}$, otherwise $x_{\text{Co}} = 0$) and this function converges to $T = \alpha T_{\text{in}}/(\alpha + f_{\text{max}})$ in the limit $x_{\text{Co}} \to \infty$. Eq (A.1b) is, on the other hand, a decreasing function of $T$. The last null-cline, Eq (A.1c) is the $T$ axis.

There exist two equilibria. One is the trivial equilibrium $(T, x) = (T_{\text{in}}, 0)$, which is an intersection of Eqs (A.1a) and (A.1c). The other is a non-trivial one $(T, x) = (T^*, x_i^*)$ where $T^*$ and $x_i^*$ are defined in Eqs (9a) and (9b). Such an equilibrium is the intersection of Eqs (A.1a) and (A.1b). To investigate the linear stability of this equilibrium, we evaluate the $2 \times 2$ Jacobian matrix:

$$J = \begin{pmatrix} -\left\{f\left(x_{\text{Co}}\right) + \alpha\right\} & -Tf'\left(x_{\text{Co}}\right) \\ -\delta_i'(T)x_i & r_i(1-2x_i) - \delta_i(T) - \alpha \end{pmatrix}. \tag{A.2}$$

An equilibrium is linearly stable if and only if the maximum value of the real part of the eigenvalue of the Jacobian matrix is negative. From the Routh-Hurwitz criteria (Murray, 2002, Appendix B), the maximum value of the real part of the eigenvalue of the $2 \times 2$ Jacobian matrix is negative if and only if the following two inequalities are satisfied:

$$\text{tr}J < 0, \tag{A.3a}$$

$$\text{det}J > 0. \tag{A.3b}$$

At the trivial equilibrium in the mono-culture of cooperators $i \in \{\text{sCo}, \text{rCo}\}$, the trace and the determinant of the Jacobian matrix are

$$\text{tr}J = r_i - \delta_i\left(T_{\text{in}}\right) - 2\alpha, \tag{A.4a}$$

$$\text{det}J = -\alpha\left\{r_i - \delta_i\left(T_{\text{in}}\right) - \alpha\right\}. \tag{A.4b}$$

If $r_i > \delta_i(T_{\text{in}}) + \alpha$, the trivial equilibrium is unstable because $\det J < 0$. On the other hand, the trivial equilibrium is stable if $r_i < \delta_i(T_{\text{in}}) + \alpha$ as the trace of the Jacobian matrix is negative and the determinant of the Jacobian matrix is positive. When the former condition holds, cooperators can invade from rare.

At the non-trivial equilibrium for cooperators $i \in \{\text{sCo}, \text{rCo}\}$, the trace of the Jacobian matrix is always negative because

$$\mathrm{tr}J = -\{f(x_i^*) + \alpha\} + \underbrace{\{r_i(1 - x_i^*) - \delta_i(T^*) - \alpha\}}_{=0 \quad \because \text{Eq (9b)}} - r_i x_i^* < 0. \tag{A.5}$$

Therefore, the non-trivial equilibrium is stable if and only if

$$\begin{aligned}
\det J &> 0 \\
\Leftrightarrow \{f(x_i^*) + \alpha\} r_i x_i^* - T^* f'(x_i^*) \delta_i'(T^*) x_i^* &> 0 \\
\Leftrightarrow r_i \{f(x_i^*) + \alpha\} &> T^* f'(x_i^*) \delta_i'(T^*) \\
\Leftrightarrow -\frac{r_i}{\delta_i'(T^*)} &< -\frac{\alpha T_{\text{in}} f'(x_i^*)}{\{\alpha + f(x_i^*)\}^2} \quad (\because T^* = \alpha T_{\text{in}} / \{\alpha + f(x_i^*)\})
\end{aligned} \tag{A.6}$$

Inequality (A.6) represents the relationship between the slope of the tangent lines of the two null-clines of Eqs (A.1a) and (A.1b) at the non-trivial equilibrium. The two tangent lines are given as below:

$$T = -\frac{\alpha T_{\text{in}} f'(x^*)}{\{\alpha + f(x^*)\}^2}(x - x^*) + T^*, \tag{A.7a}$$

$$T = -\frac{r_i}{\delta_i'(T^*)}(x - x^*) + T^*. \tag{A.7b}$$

If inequality (A.6) is satisfied, the null-cline Eq (A.1a) should exist under (above) the null-cline Eq (A.1b) at $x_i = x_i^* + \epsilon$ (or, $x_i = x_i^* - \epsilon$) where $0 < \epsilon \ll 1$.

When $r_i > \delta_i(T_{\text{in}}) + \alpha$ is satisfied, there exists at least one stable and non-trivial equilibrium point. In a mono-culture of cooperators, the null-cline Eq (A.1a) is a decreasing function of the density of the cooperators $x_i$ and converges to $\alpha T_{\text{in}} / (\alpha + f_{\text{max}})$ in the limit of $x_i \to \infty$ while the null-cline Eq (A.1b) is a decreasing function of the toxic concentration $T$ and converges to $1 - \alpha / r_i$ in the limit of $T \to 0$. Notice that $r_i > \alpha$ is a necessary condition for $r_i > \delta_i(T_{\text{in}}) + \alpha$. When $r_i > \delta_i(T_{\text{in}}) + \alpha$ is satisfied, $0 < 1 - \alpha / r_i < 1$, and therefore, there exists at least one intersection of the two null-clines. If a unique intersection exists, this point is a stable and non-trivial equilibrium point because the null-cline Eq (A.1b) exists above the null-cline Eq (A.1a) within $x \in [0, x_i^*)$ (Fig.A.1 left). If instead multiple intersections exist, all those that satisfy inequality (A.6) are stable and the dynamics from $(x_0, T_{\text{in}})$ converge to the intersection with the smallest value of $x_i^*$. It should be noted that when $r_i < \delta_i(T_{\text{in}}) + \alpha$, the trivial equilibrium is stable, and the dynamics from $(x_0, T_{\text{in}})$ converge to it, even if a second stable non-trivial

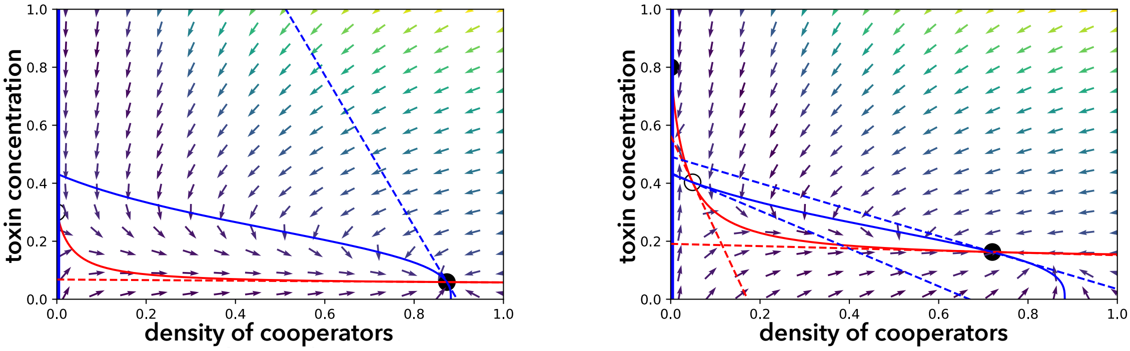


Figure A.1: Phase plane analysis

Two examples of the phase plane analysis of a mono-culture of sensitive cooperators are shown ($i = \text{sCo}$). The red solid line represents the null-clines defined by Eq (A.1a) and the two blue solid lines show the null-clines of Eqs (A.1b) and (A.1c). The dashed lines represent the tangents of the null-clines at the non-trivial equilibria given by Eqs (A.7a) and (A.7b), respectively. Left: When $r_i > \delta_i \left(T_{\text{in}}\right) + \alpha$, the dynamics converge to the non-trivial stable equilibrium (black circle). Right: When $r_i < \delta_i \left(T_{\text{in}}\right) + \alpha$, the non-trivial equilibrium closest to the trivial equilibrium is unstable (white circle), although there exists another non-trivial equilibrium that is stable. Notice that the trivial equilibrium $\left(T_{\text{in}}, 0\right)$ is unstable if $r_i > \delta_i \left(T_{\text{in}}\right) + \alpha$ (left) but stable if $r_i < \delta_i \left(T_{\text{in}}\right) + \alpha$ (right). The parameter values are: left $T_{\text{in}} = 0.3$, $n = 3$, $c_d = 0.15$, $r = 1$, $K_d = 0.2$, $K_s = 0.3$, $d_{\text{max}} = 1.0$, $f_{\text{max}} = 0.5$, and $\alpha = 0.1$; right $T_0 = 0.8$ and the other parameter values are the same as in the left panel. Notice that in the case of the mono-culture of resistant cooperators, the parameters should be changed as follows: $K_s \to K_r$ and $c_d \to c_d + c_r$.

equilibrium exists (Fig. A.1 right). In summary, cooperators can grow and the population dynamics converge to a non-trivial equilibrium if and only if and only if $r_i > \delta_i \left(T_{\text{in}}\right) + \alpha$ where $i \in \{\text{sCo}, \text{rCo}\}$.

In the case of a mono-culture of cheaters, the null-cline Eq (A.1a) is a line $T = T_{\text{in}}$. If cooperator $i$ satisfies $r_i > \delta \left(T_{\text{in}}\right) + \alpha$, there exist a unique stable equilibrium for cheaters that have the same level of resistance as the cooperators, because the cheaters' intrinsic growth rate is larger than that of the cooperators. To sum up, we obtain inequality (8) in the main text.

### Appendix 2 Conditions where cooperators invade cheaters

Here, we analyze the conditions where one strategy $i$ can invade a population of strategy $j$. It is assumed here that the population of strategy $j$ converges to the mono-culture equilibrium state prior to invasion (i.e., $(x_j, T) = \left(x_j^*, T_j^*\right)$). From Eq (9b), the following equation is satisfied:

$$r_j \left(1 - x_j^*\right) - \delta_j \left(T_j^*\right) - \alpha = 0. \tag{A.8}$$

Now, let us assume that a small number of cells in the population mutate into strategy $i$. Then, strategy $i$ can invade the population if and only if

$$r_i \left(1 - x_j^*\right) - \delta_i \left(T_j^*\right) - \alpha > 0, \tag{A.9}$$

because the total cell density is still $x_j^*$.

For ease of following the analysis, let us define $W_i(T)$ as shown in Eq (2) in the main text. Note that strategy $i$ can grow at toxin concentration $T$ if $W_i(T) > 1/\left(1-\sum_j x_j\right)$. However, $W_i(T) > W_j(T)$ implies that strategy $i$ can increase faster or decrease slower than strategy $j$. From Eq (A.8) and inequality (A.9), strategy $i$ invades the population of strategy $j$ if and only if

$$W_i\left(T_j^*\right) > W_j\left(T_j^*\right). \tag{A.10}$$

Note that cooperators cannot invade a population of cheaters which have the same level of resistance $(i,j) \in \{(\mathrm{sCo}, \mathrm{sCh}), (\mathrm{rCo}, \mathrm{rCh})\}$ because cooperators have a lower growth rate than cheaters due to the cost of cooperation ($r_i < r_j$), while the death rates are the same at any toxin concentration ($\delta_i(T) = \delta_j(T)$):

$$W_i(T) < W_j(T), (i,j) \in \{(\mathrm{sCo}, \mathrm{sCh}), (\mathrm{rCo}, \mathrm{rCh})\}. \tag{A.11}$$

For the same reason, cheaters exclude a population of cooperators that have the same level of resistance once the cheaters appear.

However, it is possible that cooperators can invade a population of cheaters with a different level of resistance at certain toxin concentrations. In addition, resistant cooperators (cheaters) can invade sensitive cooperators (cheaters), respectively, and vice versa. For such cases, inequality (A.10) is simplified using Eqs (3) and (4) in the main text as follows:

$$\begin{aligned}
& W_i\left(T_j^*\right) > W_i\left(T_j^*\right) \\
\Leftrightarrow\ & \frac{r_i}{\delta_i\left(T_j^*\right)+\alpha} > \frac{r_j}{\delta_j\left(T_j^*\right)+\alpha} \\
\Leftrightarrow\ & \frac{r_i}{d_{\max}\frac{T_j^{*n}}{T_j^{*n}+K_i^n}+\alpha} > \frac{r_j}{d_{\max}\frac{T_j^{*n}}{T_j^{*n}+K_j^n}+\alpha} \\
\Leftrightarrow\ & \alpha\left(r_i-r_j\right)\left(\tau_j^*+K_i^n\right)\left(\tau_j^*+K_j^n\right)+d_{\max}\left\{r_i\tau_j^*\left(\tau_j^*+K_i^n\right)-r_j\tau_j^*\left(\tau_j^*+K_j^n\right)\right\} > 0 \\
\Leftrightarrow\ & F_{ij}\left(\tau=\tau_j^*\right) \equiv \left(\alpha+d_{\max}\right)\left(c_j-c_i\right)\tau_j^{*2}+\left\{d_{\max}(1-c_i)K_i^n-d_{\max}(1-c_j)K_j^n+\alpha(c_j-c_i)(K_i^n+K_j^n)\right\}\tau_j^* \\
& +\alpha(c_j-c_i)K_i^nK_j^n > 0,
\end{aligned} \tag{A.12}$$

where $\tau = T^n$, and $c_i$ and $c_j$ represent the total cost of strategies $i$ and $j$, respectively. Note that as $F_{ij}(\tau)$ is a quadratic function of $\tau$, one can analytically find the range of $\tau_j^*$ where $F_{ij}\left(\tau=\tau_j^*\right)$ is satisfied once the parameter values used in inequality (A.12) have been obtained (Fig. A.2).

At the point when a small number of resistant (sensitive) cooperators invade a population of sensitive (resistant) cheaters ($(i,j) \in \{(\mathrm{rCo}, \mathrm{sCh}), (\mathrm{sCo}, \mathrm{rCh})\}$), the toxin concentration is close to that which is flowing into the chemostat ($T_j^* = T_{\mathrm{in}}$). In these two cases, the conditions where cooperators successfully invade cheaters with a different level of resistance at a certain toxin level are analytically derived. First,

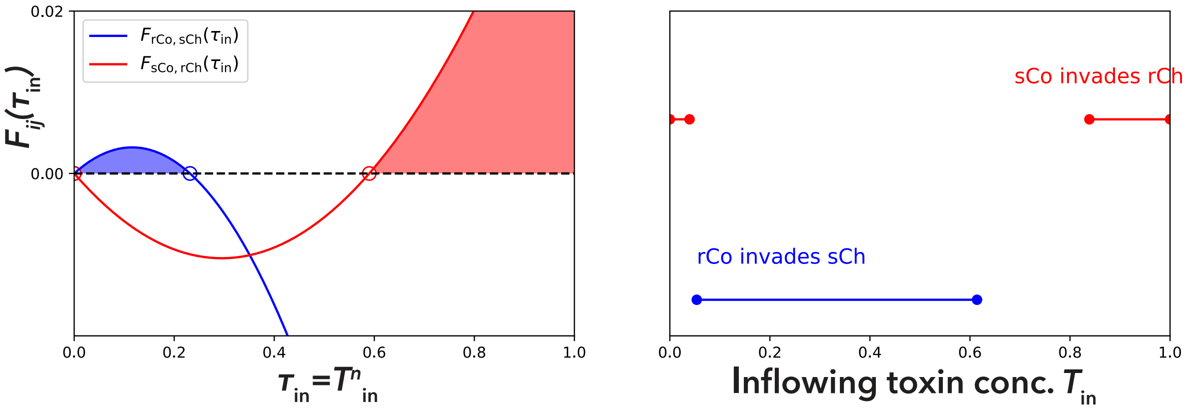


Figure A.2: Toxin concentration range where invasion of cooperators into a population of cheaters succeeds.

The range of $\tau_{\text{in}}$ (left) or $T_{\text{in}}$ (right) where the invasion of resistant cooperators into a population of sensitive cheaters (blue) or the invasion of sensitive cooperators into a population of resistant cheaters (red) succeeds. In the left panel, one can analytically find the roots of $F_{ij}(\tau_{\text{in}})$ because $F_{ij}(\tau_{\text{in}})$ is a quadratic function. The shaded blue or red area corresponds to the value of $\tau_{\text{in}}$ where invasion succeeds. In the right panel, the ranges of $T_{\text{in}}$ where the invasion succeeds in the left panel are shown; at low or high toxic concentrations, resistant cheaters are invaded by sensitive cooperators. On the other hand, at intermediate toxic concentrations, sensitive cheaters are invaded by resistant cooperators. Parameter values are $c_d = 0.1$, $c_r = 0.3$, $K_r = 0.6$, $K_s = 0.1$, $n = 3$, $\alpha = 0.1$, and $d_{\max} = 0.5$.

it is necessary to derive the range of toxin concentrations where cheaters can be maintained in mono-culture. In other words, we need to derive the range of $T_{\text{in}}$ where the density of cheaters $j$ is positive at an equilibrium state. This condition is derived by starting from the population equilibrium $x_j^*$ and solving the inequality for $T_{\text{in}}$:

$$
\begin{aligned}
& x_j^* = 1 - \frac{\delta_j(T_{\text{in}}) + \alpha}{r_j} > 0 \\
\Leftrightarrow\ & r_j > \delta_j(T_{\text{in}}) + \alpha \\
\Leftrightarrow\ & (d\text{max} - a_j)\,\tau_{\text{in}} < a_j K_j^n \\
\Leftrightarrow\ & \begin{cases} \tau_{\text{in}} \leq 1 & d\text{max} - a_j < a_j K_j^n \\ \tau_{\text{in}} < a_j K_j^n / (d\text{max} - a_j) & \text{otherwise} \end{cases}
\end{aligned}
\tag{A.13}
$$

where $\tau_{\text{in}} = T_{\text{in}}^n$ and $a_j = r_j - \alpha$. Note that $a_j > 0$ is a necessary condition that strategy $j$ can grow in mono-culture. Here, the maximum value of $\tau_{\text{in}}$ is denoted by $\hat{\tau}_{\text{in}}$:

$$
\hat{\tau}_{\text{in}} = \begin{cases} 1 & \text{if } d_{\max} - a_j < a_j K_j^n \\ a_j K_j^n / (d_{\max} - a_j) & \text{otherwise} \end{cases}
\tag{A.14}
$$

When resistant cooperators can invade sensitive cheaters, the quadratic function $F_{\text{rCosCh}}(\tau_{\text{in}})$ should be positive:

$$
\begin{aligned}
F_{\mathrm{rCosCh}}\left(\tau_{\mathrm{in}}\right) &= -\left(\alpha+d_{\max}\right) c\tau_{\mathrm{in}}^{2}+\left\{d_{\max}\left(1-c\right) K_{r}^{n}-d_{\max} K_{s}^{n}-\alpha c\left(K_{s}^{n}+K_{r}^{n}\right)\right\} \tau_{\mathrm{in}} \\
&\quad -\alpha c K_{s}^{n} K_{r}^{n}>0 \qquad (A.15)\\
\Leftrightarrow F_{1}\left(\tau_{\mathrm{in}}\right) &\equiv -F_{\mathrm{rCosCh}}\left(\tau_{\mathrm{in}}\right)<0 \qquad (A.16)
\end{aligned}
$$

where $c=c_{d}+c_{r}$. The condition that $F_{1}\left(\tau_{\mathrm{in}}\right)$ is negative for different values of $\tau_{\mathrm{in}}=[0., \hat{\tau}_{\mathrm{in}}]$ is either (i) the existence of one real root of $F_{1}\left(\tau\right)$ within $\tau_{\mathrm{in}}=[0, .\hat{\tau}_{\mathrm{in}}]$

$$
F_{1}(\hat{\tau}_{\mathrm{in}})<0 \qquad (A.17)
$$

or (ii) the existence of two real roots of $F_{1}\left(\tau_{\mathrm{in}}\right)$ (Eq (A.15)) within $\tau_{\mathrm{in}}=[0, \hat{\tau}_{\mathrm{in}}]$

$$
\left\{
\begin{array}{c}
F_{1}\left(\hat{\tau}_{\mathrm{in}}\right) \geq 0, \\
D \equiv \left\{d_{\max}(1-c)K_{r}^{n}-d_{\max}K_{s}^{n}-\alpha c(K_{r}^{n}+K_{s}^{n})\right\}^{2}-4(d_{\max}+\alpha)\alpha(K_{r}K_{s})^{n}c^{2}>0 \\
\hat{\tau}_{\mathrm{in}}>\frac{1}{2(d_{\max}+\alpha)c}\left\{d_{\max}(1-c)K_{r}^{n}-d_{\max}K_{s}^{n}-\alpha c(K_{r}^{n}+K_{s}^{n})\right\}>0,
\end{array}
\right. \qquad (A.18)
$$

because $F_{1}\left(0\right)>0$. Note that the second and third inequalities of inequalities (A.18) represents the conditions where there exist two real roots of $F_{1}\left(\tau\right)$, and where the axis of symmetry exists within $\tau_{\mathrm{in}}=[0, \hat{\tau}_{\mathrm{in}}]$, respectively. Inequality (A.17) is simplified as below:

$$
\begin{aligned}
F_{1}(\hat{\tau}_{\mathrm{in}}) &< 0 \\
\Leftrightarrow c &< \frac{d_{\max}\left(K_{r}^{n}-K_{s}^{n}\right)\hat{\tau}_{\mathrm{in}}}{(d_{\max}+\alpha)\hat{\tau}_{\mathrm{in}}^{2}+\left\{dK_{r}^{n}+\alpha(K_{r}^{n}+K_{s}^{n})\right\}\hat{\tau}_{\mathrm{in}}+\alpha K_{r}^{n}K_{s}^{n}}. \qquad (A.19)
\end{aligned}
$$

Inequalities (A.18) are, on the other hand, rewritten as below:

$$
\begin{aligned}
& F_{1}(\hat{\tau}_{\mathrm{in}}) \geq 0 \\
\Leftrightarrow\quad & c \geq \frac{d_{\max}\left(K_{r}^{n}-K_{s}^{n}\right)\hat{\tau}_{\mathrm{in}}}{(d_{\max}+\alpha)\hat{\tau}_{\mathrm{in}}^{2}+\left\{d_{\max}K_{r}^{n}+\alpha(K_{r}^{n}+K_{s}^{n})\right\}\hat{\tau}_{\mathrm{in}}+\alpha K_{r}^{n}K_{s}^{n}}, \qquad (A.20a)\\
& \hat{\tau}_{\mathrm{in}}>\frac{1}{2(d_{\max}+\alpha)c}\left\{d_{\max}(1-c)K_{r}^{n}-d_{\max}K_{s}^{n}-\alpha c(K_{r}^{n}+K_{s}^{n})\right\}>0 \\
\Leftrightarrow\quad & \frac{d_{\max}\left(K_{r}^{n}-K_{s}^{n}\right)}{d_{\max}K_{r}^{n}+\alpha\left(K_{r}^{n}+K_{s}^{n}\right)}>c>\frac{d_{\max}\left(K_{r}^{n}-K_{s}^{n}\right)}{2(d_{\max}+\alpha)\hat{\tau}_{\mathrm{in}}+d_{\max}K_{r}^{n}+\alpha\left(K_{r}^{n}+K_{s}^{n}\right)}, \qquad (A.20b)\\
& D>0 \\
\Leftrightarrow\quad & \left\{
\begin{array}{c}
c<\frac{d_{\max}(K_{r}^{n}-K_{s}^{n})}{d_{\max}K_{r}^{n}+\alpha(K_{r}^{n}+K_{s}^{n})+2\left\{\alpha(\alpha+d_{\max})K_{r}^{n}K_{s}^{n}\right\}^{1/2}} \quad \text{or} \\
(d_{\max}-c)K_{r}^{n}-d_{\max}K_{s}^{n}-\alpha c(K_{r}^{n}+K_{s}^{n})<-2\left\{(1+\alpha)\alpha\right\}^{1/2}(K_{r}K_{s})^{n/2}c<0.
\end{array}
\right. \qquad (A.20c)
\end{aligned}
$$

Note that one can ignore the case of $(d_{\max}-c)K_{r}^{n}-d_{\max}K_{s}^{n}-\alpha c(K_{r}^{n}+K_{s}^{n})<0$ when analyzing $D>0$

because of the left-hand side of inequality (A.20b). Then, inequalities (A.18) are summarized as follows:

$$\frac{d_{\max}\left(K_r^n-K_s^n\right)}{A+2\{\alpha(\alpha+1)K_r^nK_s^n\}^{1/2}}>c\geq\max\left\{\frac{d_{\max}\left(K_r^n-K_s^n\right)}{(d_{\max}+\alpha)\hat{\tau}_{\text{in}}+A+\alpha K_r^nK_s^n/\hat{\tau}_{\text{in}}},\frac{d_{\max}\left(K_r^n-K_s^n\right)}{2(d_{\max}+\alpha)\hat{\tau}_{\max}+A}\right\}\tag{A.21}$$

where $A=d_{\max}K_r^n+\alpha\left(K_r^n+K_s^n\right)$. It should be noted that

$$\begin{aligned}\frac{d_{\max}\left(K_r^n-K_s^n\right)}{(d_{\max}+\alpha)\hat{\tau}_{\text{in}}+A+\alpha K_r^nK_s^n/\hat{\tau}_{\text{in}}}&\lesseqgtr\frac{d_{\max}\left(K_r^n-K_s^n\right)}{2(d_{\max}+\alpha)\hat{\tau}_{\text{in}}+A}\\ \Leftrightarrow\frac{\alpha K_r^nK_s^n}{d_{\max}+\alpha}&\gtreqless\hat{\tau}_{\text{in}}^2\\ \Leftrightarrow\left(\frac{\alpha K_r^nK_s^n}{d_{\max}+\alpha}\right)^{1/2}&\gtreqless\hat{\tau}_{\text{in}}\quad(\because\hat{\tau}_{\text{in}}>0)\end{aligned}\tag{A.22}$$

Especially when $(\alpha K_r^nK_s^n/(d_{\max}+\alpha))^{1/2}<\hat{\tau}_{\text{in}}$, the condition where resistant cooperators can invade a population of sensitive cheaters is simplified because inequalities (A.19) and (A.21) are combined:

$$\frac{d_{\max}\left(K_r^n-K_s^n\right)}{A+2\{\alpha(\alpha+1)K_r^nK_s^n\}^{1/2}}>c.\tag{A.23}$$

When sensitive cooperators invade a population of resistant cheaters ($(i,j)=(\text{sCO},\text{rCh})$), on the other hand, the sign of $c_j-c_i=c_r-c_d=\Delta c$ determines the shape of $F_{\text{sCorCh}}(\tau)$. Sensitive cooperators can invade a population of resistant cheaters if and only if

$$\begin{aligned}F_{\text{sCorCh}}\left(\tau_{\text{in}}\right)=\;&(d_{\max}+\alpha)\Delta c\tau_{\text{in}}^2\\&+\left\{d_{\max}(1-c_d)K_s^n-d_{\max}(1-c_r)K_r^n+\alpha\Delta c\left(K_r^n+K_s^n\right)\right\}\tau_{\text{in}}\\&+\alpha\Delta cK_r^nK_s^n>0.\end{aligned}\tag{A.24}$$

When $\Delta c=0$, this inequality is simplified as $K_s^n>K_r^n$, which never holds. When instead $\Delta c>0$, sensitive cooperators can invade resistant cheaters at least at a low toxin concentration because $F_{\text{sCorCh}}(0)>0$. However, one would be interested in whether sensitive cooperators can invade resistant cheaters at a high toxin concentration. The necessary and sufficient condition for the successful invasion at a high toxin concentration is given by

$$\begin{aligned}F_{\text{sCorCh}}\left(\hat{\tau}_{\text{in}}\right)&>0\\ \Leftrightarrow\Delta c\underbrace{\left\{(d_{\max}+\alpha)\hat{\tau}_{\text{in}}+\alpha\left(K_r^n+K_s^n\right)+\alpha K_r^nK_s^n/\hat{\tau}_{\text{in}}\right\}}_{=B(\hat{\tau}_{\text{in}})}+d_{\max}K_r^nc_r+d_{\max}\left(K_s^n-K_r^n\right)&>d_{\max}K_s^nc_d\\ \Leftrightarrow\left\{B\left(\hat{\tau}_{\text{in}}\right)+d_{\max}K_r^n\right\}c_r+d_{\max}\left(K_s^n-K_r^n\right)&>\left(B+d_{\max}K_s^n\right)c_d\\ \Leftrightarrow\frac{\left\{B\left(\hat{\tau}_{\text{in}}\right)+d_{\max}K_r^n\right\}c_r+d_{\max}\left(K_s^n-K_r^n\right)}{B\left(\hat{\tau}_{\text{in}}\right)+d_{\max}K_s^n}&>c_d.\end{aligned}\tag{A.25}$$

Note that $\hat{\tau}_{\text{in}}$ is a function of $c_r$ in this case as $r_j=r(1-c_r)$. If $\Delta c<0\Leftrightarrow c_r<c_d$, sensitive cooperators

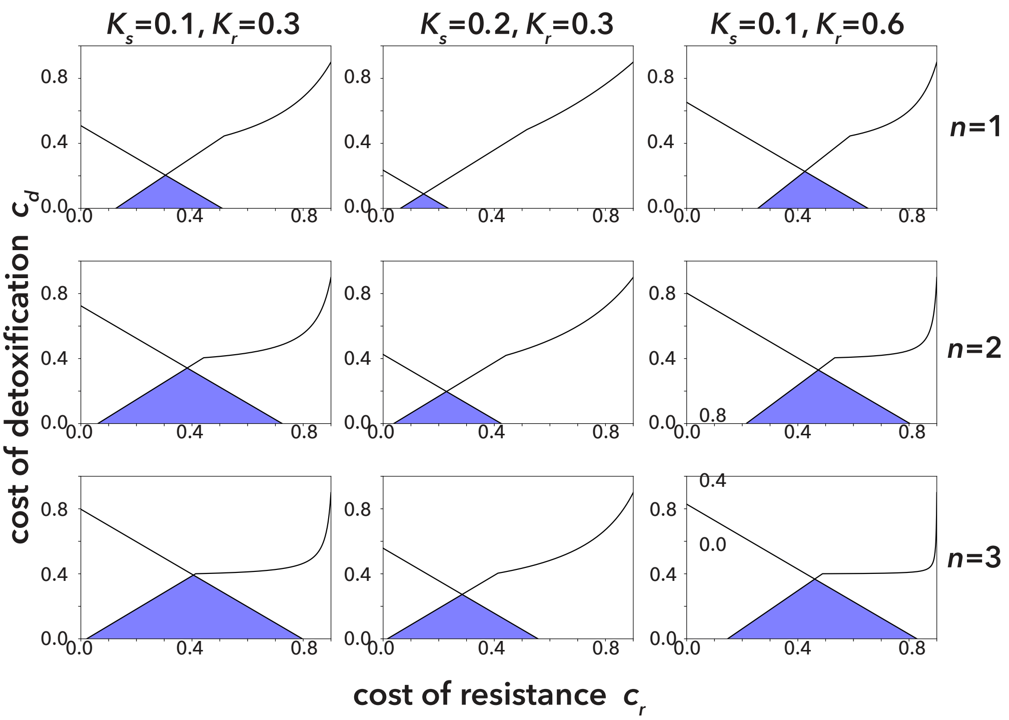


Figure A.3: The regions of the two costs where the two cooperators can invade the cheaters.

In each panel, the blue triangle area represents the region of $c_r$ and $c_d$ where the invasion of resistant cooperators to the sensitive cheaters, and the invasion of the sensitive cooperators to the resistant cheaters are successful. In the fist, second, and the third column, $(K_s, K_r) = (0.1, 0.3), (0.2, 0.3)$, and $(0.1, 0.3)$, respectively. In the first, second, and third row, $n = 1, 2$, and 3, respectively. The value of $d_{\max}$ is fixed as $d_{\max} = 0.5$ in all panels.

cannot invade resistant cheaters at any toxin concentration. In this case, $F_{\mathrm{sCorCh}}\left(\tau_0\right)$ is an upper convex function and $F_{\mathrm{sCorCh}}(0) < 0$. In addition, the axis of the symmetry of $F_{\mathrm{sCorCh}}\left(\tau_{\mathrm{in}}\right)$ is negative as below:

$$\begin{aligned} c_r &< c_d \\ \Rightarrow (1-c_d)K_s^n &< (1-c_r)K_r^n \quad (\because K_s < K_r) \\ \Rightarrow d_{\max}(1-c_d)K_s^n - d_{\max}(1-c_r)K_r^n + \alpha\Delta c\left(K_r^n + K_s^n\right) &< 0 \quad (\because \Delta c < 0). \end{aligned} \tag{A.26}$$

In other words, $F_{\mathrm{sCorCh}}\left(\tau_0\right)$ is a decreasing function of $\tau$ within $\tau = [0., \hat{\tau}_0]$ when $c_r < c_d$. In summary, sensitive cooperators can invade resistant cheaters when both $c_r > c_d$ and inequality (A.25) hold. Fig. A.3 shows the region of $c_r$ and $c_d$ and toxin concentrations where resistant cooperators can invade sensitive cheaters and sensitive cooperators can invade resistant cheaters given the values of $K_r$, $K_s$, $n$ and $d_{\max}$.

Of course, it is possible to analyze the condition where resistant cheaters can invade sensitive cheaters, and vice versa. The resistant (sensitive) cheaters invade the sensitive (resistant) cheaters when

$$W_{\mathrm{rCh}}\left(T_{\mathrm{in}}\right) \gtrless W_{\mathrm{sCh}}\left(T_{\mathrm{in}}\right) \tag{A.27}$$

holds, respectively. As $F_{\mathrm{rChsCh}}$ is a quadratic function, one can analytically find the range of $\tau_{\mathrm{in}}$ where the invasion is successful once the parameter values are obtained. When resistant (sensitive) cooperators invade the sensitive (resistant) cooperators, on the other hand, it should be noted that it is impossible to analytically calculate the value of the toxin concentration at which invasion occurs ($T_j^*$). Once this toxin concentration when invasion occurs is numerically calculated by finding the equilibrium in the mono-culture, however, one can find whether the invasion is successful or not using inequality (A.12).

### Appendix 3 Stability of the equilibria in evolutionary dynamics

As shown in Appendix 2, there can exist a range of toxin concentrations where resistant (sensitive) cooperators can invade a population of sensitive (resistant) cheaters, respectively. After the successful invasion, however, it is unclear whether resistant (sensitive) cooperators sweep out sensitive (resistant) cheaters or they coexist because the invasion of cooperators decreases the toxin concentration, thereby decreasing the death rate of each strategy. In this section, we fist show the equilibrium where two strategies $i$ and $j$ coexist. Then, we derive the conditions for which the equilibria are stable and the invader $i$ sweeps out the resident strategy $j$ or where these two strategies coexist. Although these analyses can be performed using arbitrary pairs of the four strategies (sCo, rCo, sCh, and rCh), we mainly focus on the pairs of cooperators and cheaters whose resistance levels are different (i.e., $(i, j) \in \{(\mathrm{rCo}, \mathrm{sCh}), (\mathrm{sCo}, \mathrm{rCh})\}$), because they are the only cases where a pair of strains can stably coexist.

When strategy $i$ coexists with strategy $j$ at toxin concentration $T_{ij}^{\dagger}$ the following equation should be satisfied:

$$\begin{aligned} & W_i\left(T_{ij}^{\dagger}\right) = W_j\left(T_{ij}^{\dagger}\right) \\ \Leftrightarrow\quad & \frac{r_i}{\delta_i\left(T_{ij}^{\dagger}\right) - \alpha} = \frac{r_j}{\delta_j\left(T_{ij}^{\dagger}\right) - \alpha} \\ \Leftrightarrow\quad & F_{ij}\left(\tau_{ij}^{\dagger}\right) = 0, \end{aligned} \tag{A.28}$$

where $\tau_{ij}^{\dagger} = T_{ij}^{\dagger n}$. In other words, $\tau_{ij}^{\dagger}$ is a real root of $F_{ij}(\tau)$. Therefore, cooperators cannot coexist with cheaters which have the same level of resistance (i.e., $(i, j) \in \{(\mathrm{sCo}, \mathrm{sCh}), (\mathrm{rCo}, \mathrm{rCh})\}$) because the fitness of the cooperator is always lower than that of the cheaters as shown in inequality (A.11). When resistant (sensitive) cooperators coexist with sensitive (resistant) cheaters (i.e., $(i, j) \in \{(\mathrm{rCo}, \mathrm{sCh}), (\mathrm{sCo}, \mathrm{rCh})\}$), the densities of cooperators $x_i^{\dagger}$ and cheaters $x_j^{\dagger}$ are uniquely determined because the density of cooperators is determined by the toxin concentration $T_{ij}^{\dagger}$ at equilibrium (by setting Eq (5) to 0 and solving for $x_i^{\dagger}$

and $x_j^\dagger$):

$$\begin{aligned} T_{ij}^\dagger &= \frac{\alpha T_\mathrm{in}}{\alpha + f\left(x_i^\dagger\right)} \\ \Leftrightarrow x_i^\dagger &= \frac{\alpha K_d (T_\mathrm{in} - T_{ij}^\dagger)}{f_\mathrm{max} T_{ij}^\dagger - \alpha (T_\mathrm{in} - T_{ij}^\dagger)}, \end{aligned} \tag{A.29a}$$

$$\begin{aligned} r_j\left(1 - x_i^\dagger - x_j^\dagger\right) &= \delta_j\left(T_{ij}^\dagger\right) + \alpha \\ \Leftrightarrow x_j^\dagger &= 1 - x_i^\dagger - \frac{\delta_j\left(T_{ij}^\dagger\right) + \alpha}{r_j} \\ &= 1 - x_i^\dagger - \frac{\delta_i\left(T_{ij}^\dagger\right) + \alpha}{r_i} \end{aligned} \tag{A.29b}$$

because $W_i\left(T_{ij}^\dagger\right) = W_j\left(T_{ij}^\dagger\right)$.

When two types of cooperators or cheaters coexist (i.e., $(i,j) \in \{(\mathrm{sCo}, \mathrm{rCo}), (\mathrm{sCh}, \mathrm{rCh})\}$), the densities of two strategies $i$ and $j$ are not uniquely determined. When two types of cooperators coexist, we can find the toxin density $T_{ij}^\dagger$ where they can coexist by calculating roots of $F_{i,j}(\tau)$. This toxin concentration is, however, determined only by the total density of cooperators, and therefore, the density of each cooperator type is not uniquely determined. When the two types of cheaters coexist, on the other hand, the toxin concentration remains $T_\mathrm{in}$. In other words, the necessary condition that two types of cheaters coexist is that the relative fitness of each type of cheater is the same. As the toxin density is not affected by the density of cheaters, the densities of the two types of cheaters cannot be determined. Notice that $0 < x_i^\dagger, x_j^\dagger < 1$ should be satisfied; otherwise the dynamics never converge to the equilibrium $\left(T^\dagger, x_i^\dagger, x_j^\dagger\right)$. These inequalities are rewritten as follows when $(i,j) \in \{(\mathrm{sCo}, \mathrm{rCo}), (\mathrm{sCh}, \mathrm{rCh})\}$:

$$\begin{aligned} & \begin{cases} 0 < x_i^\dagger < 1 \\ 0 < x_j^\dagger < 1 \end{cases} \\ \Leftrightarrow \quad & 0 < x_i^\dagger < 1 - \frac{\delta_j\left(T_{ij}^\dagger\right) + \alpha}{r_j} \\ \Leftrightarrow \quad & \begin{cases} 0 < x_i^\dagger \Leftrightarrow \frac{\alpha T_\mathrm{in}}{\alpha + f_\mathrm{max}} < T_{ij}^\dagger < T_\mathrm{in} \\ 0 < x_j^\dagger \end{cases} \end{aligned} \tag{A.30}$$

When strategy $i$ can invade the population of strategy $j$ at toxin concentration $T = T_j^*$, there exist two types of equilibria. One is where strategy $i$ excludes strategy $j$; i.e., $(T, x_i, x_j) = (T_i^*, x_i^*, 0)$ where $(T_i^*, x_1^*)$ represents the stable equilibrium in the mono-culture of strategy $i$. The other is where the two strategies coexist; i.e., $(T, x_i, x_j) = \left(T_{ij}^\dagger, x_i^\dagger, x_j^\dagger\right)$, where $x_i^\dagger, x_j^\dagger$ is uniquely determined when $(i,j) \in \{(\mathrm{rCo}, \mathrm{sCh}), (\mathrm{sCo}, \mathrm{rCh})\}$. Although there exist at most two values of $T^\dagger$ in each pair of $i, j$ as $F_{ij}(\tau)$ is a quadratic function, we shall find that only one value of $T^\dagger$ can provide a stable equilibrium.

To analyze the linear stability of each equilibrium, we evaluate $3 \times 3$ Jacobian matrix given as below:

$$J = \begin{pmatrix} -\{\alpha + f(x_{\mathrm{Co}})\} & -T\partial f(x_{\mathrm{Co}})/\partial x_i & -T\partial f(x_{\mathrm{Co}})/\partial x_j \\ -x_i\delta_i'(T) & r_i(1-2x_i-x_j)-\delta_i(T)-\alpha & -r_i x_i \\ -x_j\delta_j'(T) & -r_j x_j & r_j(1-x_i-2x_j)-\delta_j(T)-\alpha \end{pmatrix} \quad \text{(A.31)}$$

where $x_{\mathrm{Co}}$ is the total density of cooperators. The partial differentiation of $f(x_{\mathrm{Co}})$ is

$$\frac{\partial f(x_{\mathrm{Co}})}{\partial x_i} = \begin{cases} f_{\max}\frac{K_d}{(x_{\mathrm{Co}}+K_d)^2} & i = \mathrm{sCo}, \mathrm{rCo} \\ 0 & \text{otherwise.} \end{cases} \quad \text{(A.32)}$$

From the Routh-Hurwizt criteria, an equilibrium is linearly stable if and only if

$$\mathrm{tr}J < 0 \quad \text{(A.33a)}$$
$$\mathrm{det}J < 0 \quad \text{(A.33b)}$$
$$\sum_{k=1}^{3} M_{kk} > 0. \quad \text{(A.33c)}$$

where $M_{kk}$ is the $(k,k)$ minor of the Jacobian matrix defined by Eq (A.31).

At the equilibrium where resistant (sensitive) cooperators exclude sensitive (resistant) cheaters $(T^*, x_i^*, 0)$, the trace, the determinant, and the minors of the Jacobian matrix are

$$\mathrm{tr}J = -\{f(x_i^*)+\alpha\} - rx_i^* + \{r_j(1-x_i^*) - \delta_j(T_i^*) - \alpha\}, \quad \text{(A.34a)}$$
$$\mathrm{det}J = \{r_j(1-x_i^*) - \delta_j(T_i^*) - \alpha\} M_{33}, \quad \text{(A.34b)}$$
$$M_{11} = -rx_i^*\{r_j(1-x_i^*) - \delta_j(T^*) - \alpha\}, \quad \text{(A.34c)}$$
$$M_{22} = -(f(x_i^*)+\alpha)\{r_j(1-x_i^*) - \delta_j(T_i^*) - \alpha\}, \quad \text{(A.34d)}$$
$$M_{33} = \{\alpha + f(x_i^*)\} r_i x_i^* - T_i^* f'(x_i^*) x_i^* \delta_i'(T^*). \quad \text{(A.34e)}$$

It should be noted that $M_{33}$ should be positive because strategy $i$ persists in mono-culture. Therefore, the determinant of the Jacobian matrix is negative if and only if

$$r_j(1-x_1^*) - \delta_j(T^*) - \alpha < 0. \quad \text{(A.35)}$$

Then, $M_{11}$ and $M_{22}$ are positive and the trace of the Jacobian is negative. In short, the necessary and sufficient condition for the linear stability of the equilibrium where only strategy $i$ exists while strategy $j$ goes extinct is given by inequality (A.35). Note that at this equilibrium, the following equation should hold:

$$r_i(1-x_i^*) - \delta_i(T_i^*) - \alpha = 0. \quad \text{(A.36)}$$

Combining with this equation, inequality (A.35) is rewritten as below:

$$\begin{aligned} W_i\left(T_i^*\right) &> W_j\left(T_i^*\right) \\ \Leftrightarrow F_{ij}\left(\tau_i^*\right) &> 0. \end{aligned} \tag{A.37}$$

In other words, if the relative fitness of strategy $i$ is larger than strategy $j$ at toxin concentration $T_i^*$, the equilibrium where strategy $i$ excludes $j$ is stable.

At an equilibrium where two strategies $i$ and $j$ coexist $\left(T_{ij}^\dagger, x_i^\dagger, x_j^\dagger\right)$, the Jacobian matrix is simplified as below:

$$J = \begin{pmatrix} -\left\{\alpha + f(x_{\mathrm{Co}}^\dagger)\right\} & -T^\dagger \partial f\left(x_{\mathrm{Co}}^\dagger\right)/\partial x_i & -T^\dagger \partial f\left(x_{\mathrm{Co}}^\dagger\right)/\partial x_j \\ -x_i^\dagger \delta_i'\left(T^\dagger\right) & -r_i x_i^\dagger & -r_i x_i^\dagger \\ -x_j^\dagger \delta_j'\left(T^\dagger\right) & -r_j x_j^\dagger & -r_j x_j^\dagger \end{pmatrix}. \tag{A.38}$$

Obviously $\mathrm{tr}J < 0$ and $M_{11} = 0$, and therefore, we should evaluate $\det J$ and $M_{22} + M_{33}$. When $(i, j) = (\mathrm{sCo}, \mathrm{rCo})$, or $(\mathrm{sCh}, \mathrm{rCh})$, $J_{12} = J_{13}$, which leads to:

$$\det J = 0. \tag{A.39}$$

In other words, at least one eigenvalue of the Jacobian matrix is zero. Therefore, the coexistence of two types or cooperators or cheaters (i.e., $(i, j) \in \{(\mathrm{sCo}, \mathrm{rCo}),\ (\mathrm{sCh}, \mathrm{rCh})\}$) is not stable because the maximum value of the real part of the eigenvalues of the Jacobian matrix is zero or positive. Indeed, the coexistence of two types of cheaters is neutrally stable because $\sum_i M_{ii} = \alpha\left(r_i x_i^\dagger + r_j x_j^\dagger\right) > 0$ and $\mathrm{tr}J < 0$, suggesting that the other two eigenvalues of the Jacobian matrix are negative. Intuitively, this is because neutral selection works when sCh coexists with rCh; when one cheater changes its resistance level by mutation, there is no force to recover the densities of each type of cheater before the mutation. The coexistence of two types of cooperators, on the other hand, is neutrally stable if $\sum_i M_{ii} > 0$; otherwise this equilibrium is unstable because a positive eigenvalue exists.

When cooperators coexist with cheaters of different resistance levels (i.e., $(i, j) \in \{(\mathrm{rCo}, \mathrm{sCh}),\ (\mathrm{sCo}, \mathrm{rCh})\}$), however, coexistence can be stable. In this case $J_{13} = 0$, and the determinant of the Jacobian matrix is

$$\begin{aligned} &\det J = -\underbrace{T^\dagger x_i^\dagger x_j^\dagger \partial f/\partial x_i}_{>0}\left\{r_i \delta_j'\left(T_{ij}^\dagger\right) - r_j \delta_i'\left(T_{ij}^\dagger\right)\right\} < 0 \\ \Leftrightarrow\quad & r_j \delta_i'\left(T_{ij}^\dagger\right) < r_i \delta_j'\left(T_{ij}^\dagger\right) \\ \Leftrightarrow\quad & r_j \left\{\delta_i(T) + \alpha\right\}'\big|_{T=T_{ij}^\dagger} < r_i \left\{\delta_j(T) + \alpha\right\}'\big|_{T=T_{ij}^\dagger} \\ \Leftrightarrow\quad & \left.\frac{dG_{ij}}{dT}\right|_{T=T_{ij}^\dagger} < 0, \end{aligned} \tag{A.40}$$

where $G_{ij}(T) = r_j\{\delta_i(T) + \alpha\} - r_i\{\delta_j(T) + \alpha\}$. Then, one can find

$$\frac{dG}{dT} = \frac{dG}{d\tau}\frac{d\tau}{dT} < 0 \tag{A.41}$$

$$\Leftrightarrow \quad \frac{dG}{d\tau} < 0 \quad \because \frac{d\tau}{dT} = nT^{n-1} > 0. \tag{A.42}$$

It should be noted that $G_{ij}(\tau)$ is a part of $F_{ij}(\tau)$ as below:

$$F_{ij}(\tau) = -G_{ij}(\tau)\left(\tau + K_i^n\right)\left(\tau + K_j^n\right)/r. \tag{A.43}$$

As $\tau_{ij}^\dagger$ is non-negative, the roots of $F_{ij}(\tau)$ (i.e., $\tau_{ij}^\dagger$) should be roots of $G_{ij}(\tau)$. Then,

$$\begin{aligned} & \left.\frac{dG_{ij}}{d\tau}\right|_{\tau=\tau_{ij}^\dagger} < 0 \\ \Leftrightarrow \quad & \left.\frac{dF_{ij}}{d\tau}\right|_{\tau=\tau^\dagger} = -\left.\frac{dG_{ij}}{d\tau}\right|_{\tau=\tau_{ij}^\dagger} \underbrace{\left(\tau_{ij}^\dagger + K_i^n\right)\left(\tau_{ij}^\dagger + K_j^n\right)/r}_{>0} \\ & + \underbrace{G\left(\tau_{ij}^\dagger\right)}_{=0}\frac{2\tau_{ij}^\dagger + K_r^n + K_s^n}{r} > 0. \end{aligned} \tag{A.44}$$

$$\therefore \quad \det J > 0 \Leftrightarrow \left.\frac{dF}{d\tau}\right|_{\tau=\tau_{ij}^\dagger} > 0. \tag{A.45}$$

As shown in Appendix 2, $F_{ij}(\tau)$ is a convex (concave) quadratic function when resistant (sensitive) cooperators coexist with sensitive (resistant) cheaters, and therefore, only the smaller (larger) root of $F_{ij}(\tau)$ satisfies inequality (A.45). From here, $\tau_{ij}^\dagger$ is assumed to represent the smaller (larger) root of $F$ when resistant (sensitive) cooperators invade a population of sensitive (resistant) cheaters.

The last necessary and sufficient condition for the stable coexistence of two strategies ($(i,j) \in \{(\mathrm{rCo}, \mathrm{sCh}), (\mathrm{sCo}, \mathrm{rCh})\}$) is that the sum of the three minors of the Jacobian matrix given by Eqs (A.34c) - (A.34e) is positive:

$$\begin{aligned} \sum_i M_{ii} > 0 \quad \Leftrightarrow \quad & \left\{\alpha + f\left(x_i^\dagger\right)\right\}\left(r_i x_i^\dagger + r_j x_j^\dagger\right) > T_{ij}^\dagger f'(x_i^\dagger) x_i \delta_i'(T_{ij}^\dagger), \\ \Leftrightarrow \quad & -\frac{r_i x_i^\dagger + r_j x_j^\dagger}{x_i^\dagger \delta_i'\left(T_{ij}^\dagger\right)} < -\frac{\alpha T_{\mathrm{in}} f'\left(x_i^\dagger\right)}{\left\{\alpha + f\left(x_i^\dagger\right)\right\}^2}, \end{aligned} \tag{A.46}$$

because $M_{11} = 0$. Notice that the sufficient condition of inequality (A.46) is:

$$-\frac{r_i}{\delta_i'\left(T_{ij}^\dagger\right)} < -\frac{\alpha T_{\mathrm{in}} f'\left(x_i^\dagger\right)}{\left\{\alpha + f\left(x_i^\dagger\right)\right\}^2} \tag{A.47}$$

because $x_j^\dagger > 0$. As we have shown in Appendix 1, inequality (A.47) represents the relationship between

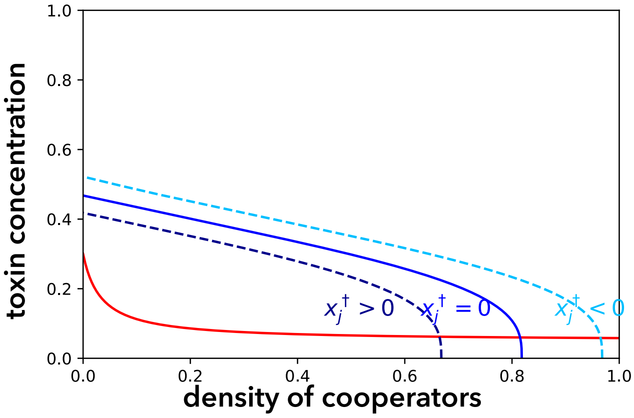


Figure A.4: Schematic illustration of the phase plane analysis in the evolutionary dynamics

Fixing $x_j = x_j^\dagger$, one can find the equilibria in $x_i - T$ phase plane. The equilibrium where cooperators exclude cheaters is the intersection of the null-clines given by Eqs (A.1a) and (A.1b) with $x_j^\dagger = 0$ (solid red and blue lines, respectively). If the equilibrium where cooperators and cheaters coexist exists, $x_2^\dagger$ should be positive (dashed navy line). In this case $T_{ij}^\dagger > T_i^*$. If $T_{ij}^\dagger < T_i^*$, on the other hand, $x_2^\dagger < 0$ (dashed sky blue line), meaning that no such equilibrium exists within $0 < T, x_i, x_j < 1$.

the slopes of the two tangent lines of the two null-clines below:

$$T = \frac{\alpha T_{\mathrm{in}}}{\alpha + f(x_i)}, \tag{A.48a}$$

$$x_i = 1 - \frac{\delta_i(T) + \alpha}{r_i} - x_j. \tag{A.48b}$$

By fixing $x_j = x_j^\dagger$, one can find that Eq (A.1b) is a special case of Eq (A.48b) when $x_j^\dagger = 0$. In other words, the null-cline given by Eq (A.48b) is obtained by parallel moving the null-cline Eq (A.1b) along the $x_i$ axis, and the intersection(s) of the two null-clines given by Eqs (A.48a) and (A.48b) is an equilibrium $\left(T^\dagger, x_j^\dagger, x_j^\dagger\right)$ (Fig. A.4). When $x_j^\dagger = 0$, on the other hand, the intersection(s) of the two null-clines represents the equilibrium of $(T_i^*, x_i^*, 0)$. It should be noted that the dynamics never converge to the equilibrium where strategies $i$ and $j$ coexist when $T_i^* > T_{ij}^\dagger \Leftrightarrow \tau_i^* > \tau_{ij}^\dagger$, because $x_2^\dagger < 0$. As $F_{\mathrm{rCo,sCh}}(\tau)$ is a convex quadratic function, the equilibrium $(T_{\mathrm{rCo}}^*, x_{\mathrm{rCo}}^*, 0)$ is stable when the equilibrium where these two strategies coexist is not feasible. In other words, there exists at most one stable equilibrium when resistant cooperators invade sensitive cheaters. When sensitive cooperators invade resistant cheaters, on the other hand, there exist at most two stable equilibria as $F_{\mathrm{sCo,rCh}}(\tau)$ is a concave quadratic function (Fig. A.5). If the two equilibria are unstable, oscillations occur (Fig. A.6). However, Monte-Carlo simulations suggest that the parameter range which causes the oscillations is very small (Fig. A.7).

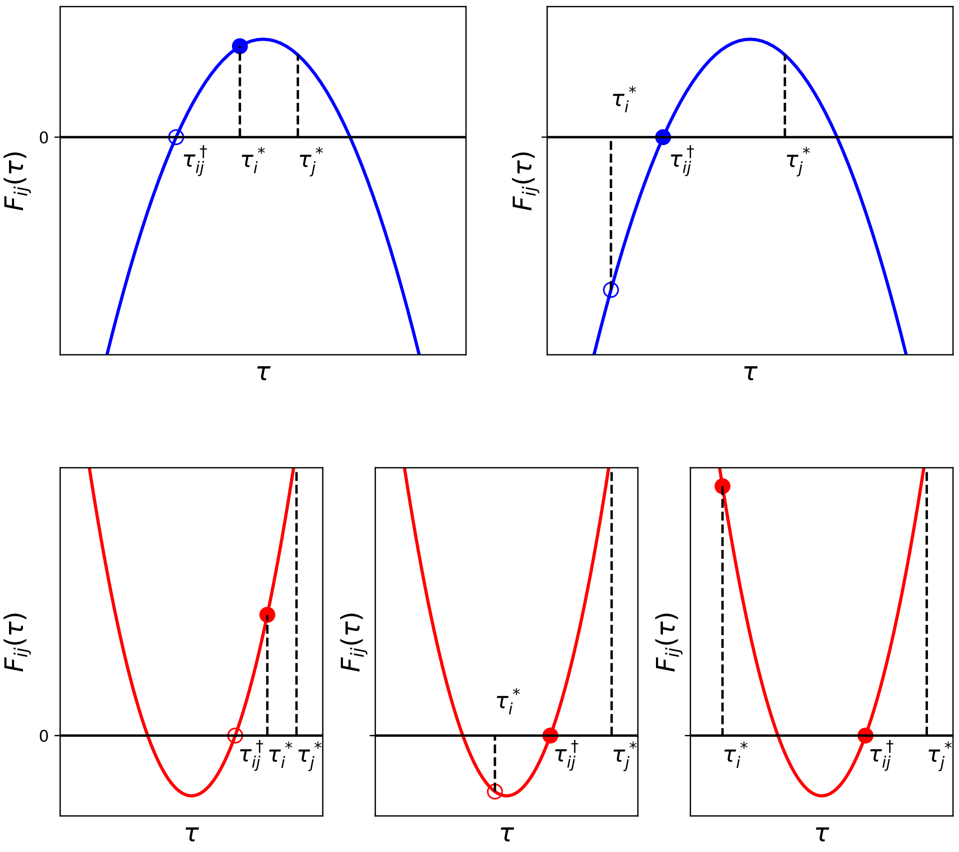


Figure A.5: Schematic illustrations of $F_{ij}(\tau)$ and the stability of two equilibria

Schematic illustrations of the quadratic function of $F_{ij}(\tau)$ and examples of two equilibria when $(i, j) = (\mathrm{rCo}, \mathrm{sCh})$ (top) or $(i, j) = (\mathrm{sCo}, \mathrm{rCh})$ (bottom) are shown. $F_{ij}\left(\tau_j^*\right) > 0$ is a necessary and sufficient condition for the invasion of strategy $i$ into the population of strategy $j$ in both cases. When $(i, j) = (\mathrm{rCo}, \mathrm{sCh})$, $F_{ij}(\tau)$ is a convex function and $\tau_{ij}^\dagger$ is the smaller root of $F_{ij}(\tau)$. If $\tau_i^* > \tau_{ij}^\dagger$ (top left), the equilibrium where the resistant cooperators exclude the sensitive cheaters is stable when $F_{ij}\left(\tau_{ij}^\dagger\right) > 0$ while the equilibrium of the coexistence does not exist as $x_j^\dagger < 0$. When $\tau_i^* < \tau_{ij}^\dagger$, on the other hand, the equilibrium where the resistant cooperators exclude the sensitive cheaters is unstable but the equilibrium of the coexistence exists. In the case of $(i, j) = (\mathrm{sCo}, \mathrm{rCh})$, $F_{ij}(\tau)$ is a concave function and $\tau^\dagger$ is the larger root of $F_{ij}(\tau)$. As in the case of $(i, j) = (\mathrm{rCo}, \mathrm{sCh})$, the equilibrium where the sensitive cooperators exclude the resistant cheaters is stable but the equilibrium of the coexistence does not exist (bottom left). However, $\tau_{ij}^\dagger > \tau_i^*$ does not mean the equilibrium of the exclusion of the resistant cheaters is unstable; when $\tau_i^*$ is larger than the smaller root of $F_{ij}(\tau)$, only the equilibrium of the coexistence can be stable (bottom center). On the other hand, if $\tau_i^*$ is smaller than the smaller root of $F_{ij}(\tau)$, the equilibrium of the exclusion is again stable, suggesting that both equilibria can be stable (bottom right).

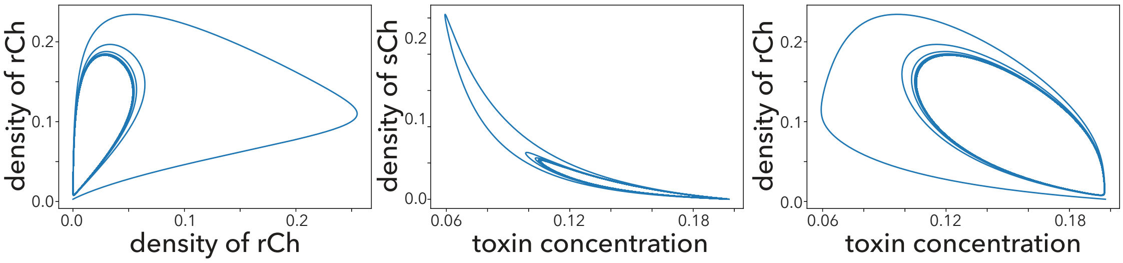


Figure A.6: Oscillation in evolutionary dynamics

Oscillation appears when the equilibria of exclusion and coexistence are unstable. These figures show an example of $(i, j) = (\mathrm{sCo}, \mathrm{rCh})$. Parameter values are $c_r = 0.672595$,, $c_d = 0.175532$, $K_s = 0.087769$, $K_r = 0.759002$, $n = 1$, $T_0 = 0.197481$, $\alpha = 0.120010$, $d_{\max} = 1$, $f_{\max} = 0.5$, $K_d = 0.2$, and $r = 1$.

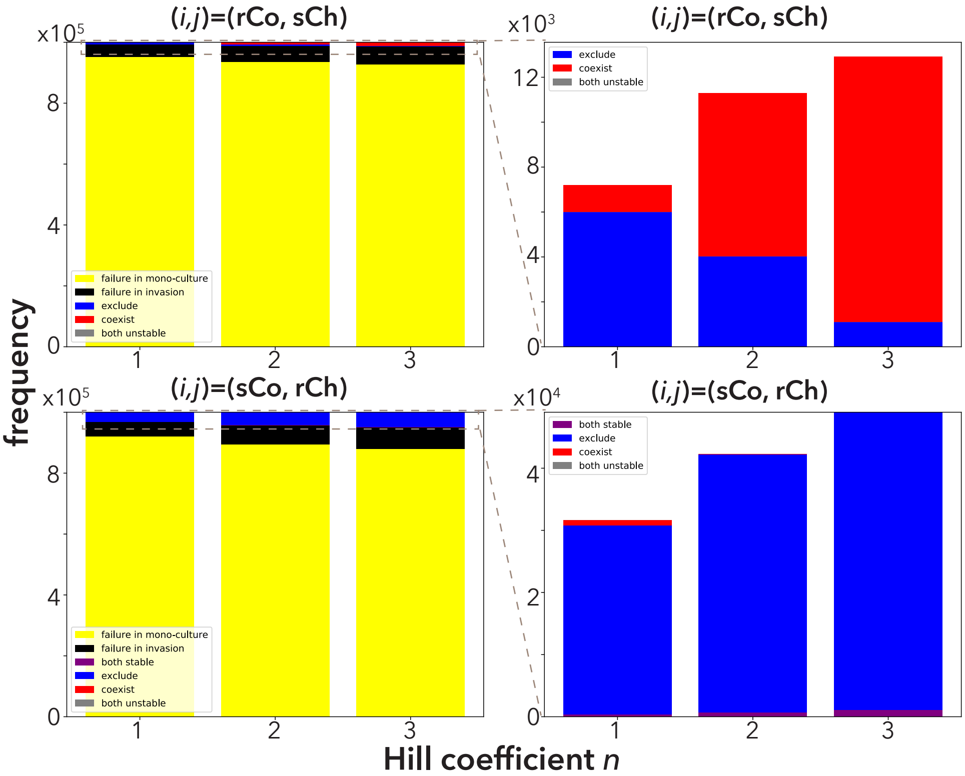


Figure A.7: Stability analysis with Monte Carlo simulation

To check whether at least one equilibrium is stable when cooperators invade cheaters, the linear stability of the equilibria of coexistence and exclusion are analyzed with a set of parameters $K_s, K_r, c_r, c_d, T_0, \alpha$ sampled using the Monte Carlo method with three values of the Hill coefficient $(n = 1, 2, 3)$. Each parameter except $K_r$ is sampled from the uniform distribution $\mathcal{U}(0, 1)$ and $K_r$ is sampled from $\mathcal{U}(K_s, 1)$. The Monte Carlo simulation sampled $10^6$ sets of parameters in each case. First row: $(i, j) = (\mathrm{rCo}, \mathrm{sCh})$. Regardless of the value of $n$, the majority of parameter sets show the failure of mono-culture of rCo and/or sCh (yellow in left), or the failure of the invasion of rCo into sCh (black in left). When rCo can invade sCh, either the equilibrium of exclusion (blue) or that of coexistence (red) is stable. In other words, there is no parameter set wherein both equilibria are unstable (gray). As $n$ gets larger, the frequency of stable exclusion (blue) becomes smaller while that of coexistence (red) becomes larger when rCo invades sCh. Second row: $(i, j) = (\mathrm{sCo}, \mathrm{rCh})$. The frequency of stable exclusion increases as $n$ grows while that of coexistence decreases. Notice that in this case, it is possible that both equilibria are stable. The fixed parameter values are $d_{\max} = 1$, $f_{\max} = 0.5$, $K_d = 0.2$, and $r = 1$.

### Appendix 4 Three-strategy situations

If two strategies $i, j$ stably coexist, mutation can provide the third strategy. Notice that the stable coexistence of the two strategies occurs only if $(i, j) \in \{(\mathrm{rCo}, \mathrm{sCh}), (\mathrm{sCo}, \mathrm{rCh})\}$. As the fitness of cooperators is lower than that of cheaters which have the same level of the resistance, these cooperators (i.e., sCo and rCo, respectively) disappear even when they arrive by mutation. In this section, we shall see the evolutionary dynamics where the new cheaters (i.e., rCh or sCh) invade the population where two strategies, rCo and sCh, or sCo and rCh coexist.

At the beginning, two strategies $i$ and $j$ ($(i, j) \in \{(\mathrm{rCo}, \mathrm{sCh}), (\mathrm{sCo}, \mathrm{rCh})\}$) coexist and the following equation should hold in Appendix 3:

$$W_i\left(T_{ij}^{\dagger}\right) = W_j\left(T_{ij}^{\dagger}\right). \tag{A.49}$$

Then, a new strategy $k$, a cheater with the same level of resistance as strategy $i$ ($(i, k) \in \{(\mathrm{sCo}, \mathrm{sCh}), (\mathrm{rCo}, \mathrm{rCh})\}$), can invade the population once it appears by the mutation. This is because

$$W_k(T) > W_i(T) \tag{A.50}$$

holds regardless the value of $T$. Hence, the two strategies $i$ and $k$ never coexist. In addition, the coexistence of two strategies $j$ and $k$ cannot be stable as the cheaters cannot change the toxin concentration (see Appendix 3). Therefore, the only stable equilibrium is the mono-culture of one of the three strategies.

Indeed, the only stable equilibrium is the mono-culture of strategy $k$. At the equilibrium of the mono-culture of strategy $k$ ($(T, x_i, x_j, x_k) = (T_{\mathrm{in}}, 0, 0, x_k^*)$), the Jacobian matrix is written as below:

$$J = \begin{pmatrix} -\alpha & -T\partial f/\partial x_i & 0 & 0 \\ 0 & r_i(1 - x_k^*) - \delta_i(T_{\mathrm{in}}) - \alpha & 0 & 0 \\ 0 & 0 & r_j(1 - x_k^*) - \delta_j(T_{\mathrm{in}}) - \alpha & 0 \\ -x_k^*\delta_k'(T_{\mathrm{in}}) & -r_k x_k^* & -r_k x_k^* & -r_k x_k^* \end{pmatrix}. \tag{A.51}$$

Then, the eigenvalues of the Jacobian matrix $\lambda$ are the diagonal elements of the Jacobian matrix by cofactor expansion:

$$\begin{aligned} & |\lambda I - J| = 0 \\ \Leftrightarrow\ & \prod_{l=1}^{4}(\lambda - J_{ll}) = 0 & \text{(A.52)} \\ \therefore\ & \lambda = J_{ll}, l = 1, 2, 3, 4. & \text{(A.53)} \end{aligned}$$

$J_{11}$ and $J_{44}$ are negative. In addition, $J_{22}$ is also negative for the following reason:

$$\begin{aligned} & W_i\left(T_{\text{in}}\right) < W_k\left(T_{\text{in}}\right) = \frac{1}{1 - x_k^*} \\ \Leftrightarrow \quad & r_i(1 - x_k^*) - \delta_i\left(T_{\text{in}}\right) - \alpha < r_k(1 - x_k^*) - \delta_k\left(T_{\text{in}}\right) - \alpha = 0. \end{aligned} \tag{A.54}$$

To evaluate the sign of $J_{33}$, it should be noted that

$$W_j\left(T_{\text{in}}\right) < W_i\left(T_{\text{in}}\right), \tag{A.55}$$

which is satisfied due to the invasion of strategy $i$ to $j$. Therefore,

$$W_j\left(T_{\text{in}}\right) < W_k\left(T_{\text{in}}\right), \tag{A.56}$$

which suggests that $J_{33} < 0$ as shown in inequality(A.54). Thus, all eigenvalues are negative and the mono-culture of strategy $k$ is stable. The stability of the mono-culture of the other two strategies is analyzed similarly. However, the mono-culture of strategy $i$ or strategy $j$ is unstable because the sign of one eigenvalue is the same as $W_k\left(T_i^*\right) - W_i\left(T_i^*\right) > 0$ or $W_k\left(T_{\text{in}}\right) - W_j\left(T_{\text{in}}\right) > 0$, respectively.

### Appendix 5 ODE including mutation rates

In the main text, we ignore mutations as shown in Eq (1). This is because it is difficult to analytically find an equilibrium and to analyze the stability of the equilibrium when mutations are taken into account. In this section, we show some examples of the dynamics when mutations are introduced into Eq (1). In this case, the dynamics of the density of strategy $i$ are defined by

$$\frac{dx_i}{dt} = \left(1 - \sum_j x_j\right) \sum_j Q_{ji} r_j x_j - \left\{\delta_i\left(T\right) + \alpha\right\} x_i, \tag{A.57}$$

where $Q_{ji}$ is the probability that strategy $j$ mutates into $i$. In our scenario, there exist two independent mutations: (i) changing cooperators to cheaters and vice versa, and (ii) changing the level of resistance. The mutation probabilities are denoted as $\mu_1$ and $\mu_2$, respectively. The mutation matrix $Q = (Q_{ji})$ is composed of $\mu_1$ and $\mu_2$ as follows:

$$Q_{ji} = \begin{cases} (1-\mu_1)(1-\mu_2) & j = i \\ \mu_1(1-\mu_2) & \text{mutated only in the detoxification ability} \\ (1-\mu_1)\mu_2 & \text{mutated only in the resistance level} \\ \mu_1\mu_2 & \text{both mutated} \end{cases} \tag{A.58}$$

At an equilibrium of this case, the density of each strategy $i$ should satisfy:

$$1 - \sum_j x_j^* = \frac{\{\delta_i(T^*) + \alpha\} x_i^*}{\sum_j r_j x_j^* Q_{ji}}, \tag{A.59}$$

where $T^* = \alpha T_{\text{in}} / \left\{\alpha + f\left(x_{\text{sCo}}^* + x_{\text{rCo}}^*\right)\right\}$. Note that $x_i^* > 0$. It is difficult to find the equilibrium state, or to analyze the stability of this equilibrium as one has to calculate a $5 \times 5$ Jacobian matrix.

However, computer simulations show the effect of mutation rates on an equilibrium. When both mutation probabilities are large ($\mu_1 = 10^{-1}, \mu_2 = 10^{-1}$), all four strategies coexist at relatively large density (Fig. A.8 top left). This is because all strategies produce the other strategies with relatively large probabilities. In other words, natural selection is weak as mutation probabilities are large.

If we reduce $\mu_2$, the majority of cells are sCo or sCh, while the densities of rCo and rCh are close to (but not equal to) zero (Fig. A.8 top right). As $\mu_2$ is small, sensitive strains rarely produce resistant strains, while cheaters produces cooperators of the same level of resistance (and vice versa). In addition, as the toxin concentration is low, sensitive strains are more advantageous than resistant strains (see Appendix 2). For these reasons, most cells are sCo and sCh.

If $\mu_1$ is small but $\mu_2$ is large (Fig. A.8 bottom left), cooperators almost go extinct. Cheaters rarely produce cooperators as $\mu_1$ is small, and therefore, cooperators are excluded due to natural selection. On the other hand, both sensitive and resistant cheaters coexist as $\mu_2$ is large.

When both mutation rates are small (Fig. A.8 bottom right), one of the four strategies dominates most of the time. At the beginning, sCo grows and and the toxin concentration decreases. Following that, sCh increases and sCo are excluded. Then, rCh invades and sCh are swept out as the toxin concentration is intermediate. Notice that rCo does not increase when sCh is dominant and the toxin concentration is increasing.

In summary, multiple strategies can coexist when both or either of the two mutation probabilities are large. As the mutation probabilities decrease, this coexistence collapses because the strength of natural selection increases. In particular, cooperators almost go extinct if $\mu_1$ is small. When both $\mu_1$ and $\mu_2$ are small, we can see similar evolutionary dynamics as in the state transition in Fig. 3.

### Appendix 6 Optimum culture conditions at the equilibrium

Defining the objective function as Eq (7), one can consider optimizing the efficiency of detoxification at an equilibrium state. As the equilibria can be classified into three types (the existence of only cheaters, coexistence of cooperators with cheaters, and a mono-culture of cooperators), we can consider the optimization problem in each equilibrium class. In this section, we shall see how to obtain the maximum detoxification efficiency at each equilibrium class by changing the dilution rate $\alpha$ and the toxin concentration flowing

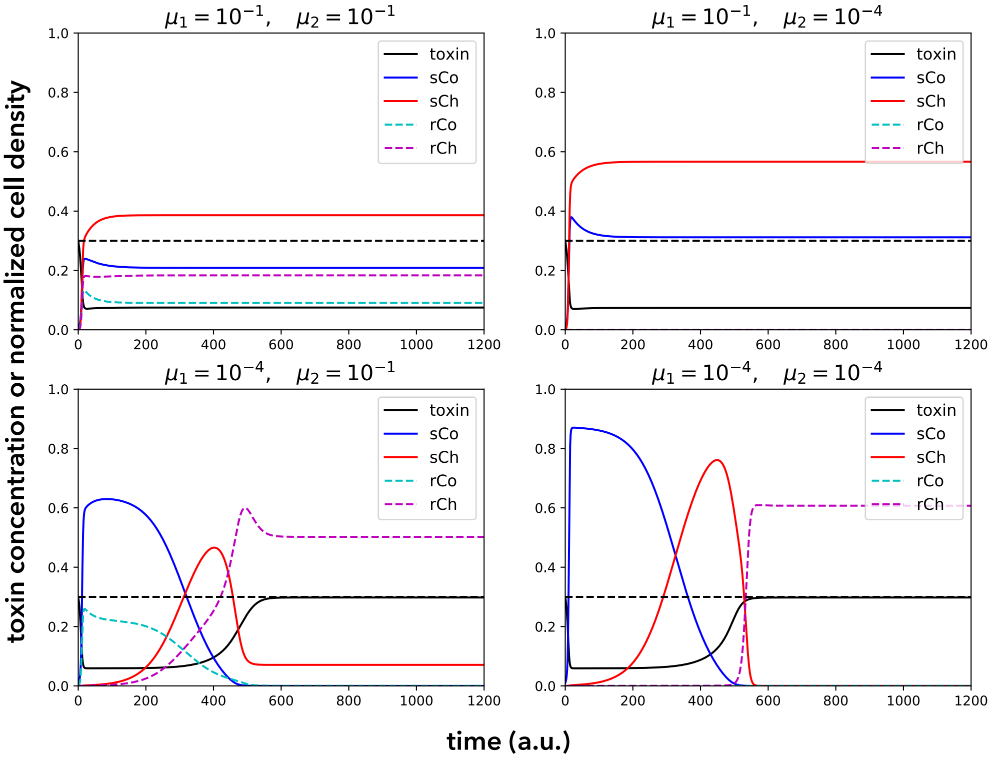


Figure A.8: Evolutionary dynamics including mutation rates in ODE

Examples of evolutionary dynamics with Eq (A.57) with different values of $\mu_1$ and $\mu_2$. In each panel, the dashed black line represents the toxin concentration flowing into the chemostat $T_{\text{in}}$, while the solid black line is the toxin concentration flowing out of the chemostat $T$. The other lines represent the densities of each strategy (solid blue: sCo, solid red: sCh, dashed cyan: rCo, and dashed magenta rCh). In each case, the initial condition is $T(0) = T_{\text{in}}$, $x_{\text{sCo}}(0) = 0.01$, and $x_i(0) = 0$ where $i \in \{\text{rCo}, \text{sCh}, \text{rCh}\}$. The other parameter values are $\alpha = 0.1$, $T_{\text{in}} = 0.3$, $f_{\text{max}} = 0.5$, $K_d = 0.2$, $r = 1$, $c_d = 0.15$, $c_r = 0.3$, $d_{\text{max}} = 1$, $K_s = 0.3$, $K_r = 0.5$, and $n = 3$.

into the system $T_{\text{in}}$.

If there exist only sensitive or resistant cheaters in the population, the toxin concentration remains that flowing into the system $T = T_{\text{in}}$, regardless of the density of cheaters. Then, the detoxification efficiency is zero.

When cooperators coexist with cheaters of different resistance level, the toxin concentration is $T^{\dagger}_{\text{rCo,sCh}}$ or $T^{\dagger}_{\text{sCo,rCh}}$. Here, we denote strategies $i$ and $j$ as cooperators and cheaters, respectively. As shown in Appendix 3, the coexistence equilibrium is analytically calculated. In this case, there exist three constraints: (i) strategy $i$ can invade the population of $j$ and strategy $j$ can persist in a mono-culture ($W_i\left(T_{\text{in}}\right) > W_j\left(T_{\text{in}}\right)$), (ii) there exists a smaller (larger) real root of $F_{ij}(\tau)$, and (iii) the coexistence equilibrium is stable, which is given by inequality (A.46):

$$
\begin{aligned}
\underset{\alpha, T_{\text{in}}}{\text{maximize}} \quad & \phi\left(\alpha, T_{\text{in}}\right) = \alpha\left(T_{\text{in}} - T^{*\dagger}_{ij}\right) \\
\text{subject to} \quad & W_i\left(T_{\text{in}}\right) > W_j\left(T_{\text{in}}\right) > 0, \\
& F_{ij}(\tau) \quad \text{has real roots}, \\
& -\left\{r_i x^{\dagger}_i + r_j x^{\dagger}_j\right\} / \left\{x^{\dagger}_i \delta'_i\left(T^{\dagger}_{ij}\right)\right\} < -\left\{\alpha T_{\text{in}} f'\left(x^{\dagger}_i\right)\right\} / \left\{\alpha + f\left(x^{\dagger}_i\right)\right\}^2.
\end{aligned}
$$

Notice that $T^{\dagger}_{ij}$ is independent of $T_{\text{in}}$. This leads to

$$\frac{\partial \phi}{\partial T_{\text{in}}} = \alpha > 0. \tag{A.60}$$

Therefore, the maximum efficiency of detoxification at this class of equilibria is given by the maximum value of $T_{\max in}$ that satisfies the constraints with each value of $\alpha$. As the constraints are complex, however, it would be easier to numerically find the maximum detoxification efficiency (Fig. 4 right column).

If only cooperators $i$ exist, the toxin concentration at equilibrium is $T^*_i$. In this case, one can consider two types of constraints: (i) the equilibrium $(T^*_i, x^*_i)$ is stable (unless mutation occurs), which is defined by inequality (A.6), or (ii) strategy $i$ can grow from low density ($r - d_i\left(T_{\text{in}}\right) - \alpha > 0$). Notice that constraint (ii) is sufficient for constraint (i). If one uses (i) as a constraint, however, the trivial equilibrium $(T, x) = (T_{\text{in},0})$ can be stable (see Fig. A.1 right), which reduces the detoxification efficiency to zero. In other words, the efficiency at equilibrium depends on the initial condition if constraint (i) is used. This problem is avoided when constraint (ii) is used, because the trivial equilibrium is unstable in this case. For this reason, we used constraint (ii) for the optimization problem with this class of equilibrium.

$$
\begin{aligned}
\underset{\alpha, T_{\text{in}}}{\text{maximize}} \quad & \phi\left(\alpha, T_{\text{in}}\right) = \alpha\left(T_{\text{in}} - T^*_i\right) \\
\text{subject to} \quad & r_i > \delta_i\left(T_{\text{in}}\right) + \alpha.
\end{aligned}
$$

Notice that it is impossible to analytically find the maximum detoxification efficiency because one cannot write $T_i^*$ in a closed form as shown in Eq (9a). However, due to the low dimensionality of the objective function, one can numerically find its global optimum (Fig. 4 left column). It should be noted that the mono-culture of cooperators can be invaded by cheaters of opposite resistance only with constraint (i) or (ii). Therefore, to achieve maximum detoxification efficiency with this class of equilibrium, it is necessary that cheaters are excluded from the population before changing the parameters $(\alpha, T_{\text{in}})$ so that the detoxification efficiency of the mono-culture of cooperators is maximized.

When cooperators can coexist with cheaters of opposite resistance with certain parameter values $(\alpha, T_{\text{in}})$, this parameter set gives the higher detoxification efficiency when there are only cooperators in the population. This is because the necessary condition for the existence of equilibria where cooperators can coexist with cheaters of opposite resistance is that $T_{ij}^{\dagger} > T_i^*$ (see Fig. A.4), although the stability of these two types of equilibria is unclear. This implies that the detoxification efficiency of a mono-culture of cooperators is larger with the parameters $(\alpha, T_{\text{in}})$ at which cooperators coexist with cheaters:

$$\phi\left(\alpha, T_{\text{in}}, T_{ij}^{\dagger}\left(\alpha, T_{\text{in}}\right)\right) < \phi\left(\alpha, T_{\text{in}}, T_i^*\left(\alpha, T_{\text{in}}\right)\right). \tag{A.61}$$

Therefore, the maximum detoxification efficiency of a mono-culture of cooperators is larger than that of a co-culture of cooperators with cheaters that differ in the level of resistance.

$$\max_{\alpha, T_{\text{in}}} \phi\left(\alpha, T_{\text{in}}, T_{ij}^{\dagger}\left(\alpha\right)\right) < \max_{\alpha, T_{\text{in}}} \phi\left(\alpha, T_{\text{in}}, T_i^*\left(\alpha, T_{\text{in}}\right)\right). \tag{A.62}$$

If cheaters can be excluded from coexistence with cooperators of different resistance by changing $(\alpha, T_{\text{in}})$, the efficiency of detoxification is maximized as follows: (i) change $(\alpha, T_{\text{in}})$ so that the equilibrium where a mono-culture of cooperators is stabilized while the equilibrium of coexistence is destabilized, (ii) maintain the parameter values until cheaters go extinct, and then, (iii) change the parameters $(\alpha, T_{\text{in}})$ so that the detoxification efficiency given by the mono-culture of cooperators is maximized. If coexistence cannot be destabilized, the maximum detoxification efficiency is achieved just by maximizing it for the co-culture of cooperators with cheaters.

### Appendix 7 Optimization of cumulative detoxification efficiency

In Appendix 2, we found that cooperators can invade a population of cheaters at some toxin concentration if the level of resistance is different. In addition, Appendix 6 shows that cooperators can exclude cheaters with different resistance level by changing the dilution rate or the toxin concentration flowing into the system, and then one can maximize detoxification efficiency in a mono-culture of cooperators. Once cheaters with the same level of resistance appear by mutation, however, cooperators are swept out and the detoxification efficiency defined by Eq (7) becomes zero as cheaters do not detoxify.

To recover detoxification, it is necessary to introduce cooperators which have a different level of resistance from the resident cheaters and to increase the toxin concentration so as to satisfy inequality (A.10), because cooperators with different resistance are unlikely to appear by mutation as they require double mutations. When the state of the population (existence of four strategies) is not observable, one problem is how often sCo and rCo should be introduced into the population in order to maximize detoxification from the beginning of the inoculation to a certain time. If cooperator inoculation probabilities are too small, the population will consist of cheaters most of the time and detoxification efficiency will be low. If cooperator inoculation probabilities are too large, cooperators are likely to be introduced into a population already containing cooperators unnecessarily, or where the invasion of cooperators fails because the resident cheaters have the same level of resistance. Such unfavorable introduction of cooperators is costly because it decreases detoxification efficiency.

If we assume a small enough mutation probability, we can redefine the time scale to discrete time steps. Then, focusing on the existence of each strategy rather than their densities, one can consider discrete population states (e.g., a mono-culture of one of the four strategies, or a transient state where two strategies coexist and one strategy is invading and sweeping out another). Combining discrete time steps and discrete population states, the model is defined as a Markov chain. We assume that only one mutation or introduction of cooperators occurs in one time step and these events never occur during the transient state. In other words, it is assumed that mutation or the introduction of cooperators occurs only when the population state is an equilibrium state of mono- or co-culture of one or two of the four strategies.

In this section, we first show the details of the special case shown in Box 1 where $\mu_2 = 0$. Note that the Markov chain remains ergodic when $\mu_2 > 0$ in this case (relaxing assumption (i)) because a positive $\mu_2$ adds transient states between sCo-rCo and sCh-rCh, respectively, and the two states where sCo (rCo) excludes sCh (rCh) or they coexist but $(\alpha, T_{\text{in}})$ are not optimized. Then, we will investigate whether relaxing the other three assumptions in Box 1 affect the ergodicity of the Markov chain or not: (ii) mutation and introduction do not occur when the population is in one of the transient states, nor at the same time, (iii) both types of cooperators can mutually exclude each other, and (iv) sCo and rCo can exclude cheaters with different resistance levels by changing $\alpha$ and $T_{\text{in}}$. In the latter part of this section, we show how to maximize the cumulative detoxification efficiency when Markov chains are not ergodic.

When $\mu_2 = 0$ and the above three assumptions in Box 1 are held, there exist at least 14 states as shown in Fig. A.9; the mono-culture of sCo (state 1), the transient state (which is unstable in short-term dynamics Eqs (1) and (5)) when sCh appears in the mono-culture of sCo by mutation (state 2), the mono-culture of sCh (state 3), the transient state when rCo is introduced to the mono-culture of sCh (state 4), the mono-culture of rCo (state 5), the transient state when rCh appears in the mono-culture of rCo by mutation (state 6), the mono-culture of rCh (state 7), the transient state when sCo is introduced to the mono-culture of rCh (state 8), the transient state when rCo is introduced to the mono-culture of sCo (state 9), the transient state when sCo is introduced to the mono-culture of rCo (state 10), the transient state when sCo is unnecessarily introduced to the mono-culture of sCo (state 11), the transient state when sCo is introduced to the mono-culture of sCh but soon excluded (state 12), the transient state when rCo is unnecessarily introduced to the mono-culture of rCo (state 13), and the transient state when rCo is introduced to the mono-culture of rCh but soon excluded (state 14). One can add more transient states between the mono-culture states when transition from one mono-culture state to another mono-culture state takes more units of time than other transitions. In addition, the number of the transition states depends on the time scale of a discrete time step. These points do not change, however, the ergodicity of the Markov chain, and we will continue the scenario with the least number of states. When $m_1, m_2 > 0$ and $m_1 + m_2 + \mu_1 < 1$, the Markov chain shown in Fig. A.9 is ergodic, and therefore, an arbitrary initial probability distribution of the states of population $\boldsymbol{\pi}(0; \boldsymbol{m}, \boldsymbol{\mu})$ converges to a unique stationary distribution $\boldsymbol{\pi}^*(\boldsymbol{m}; \boldsymbol{\mu})$, which is obtained below:

$$\sum_{i=1}^{14} \pi_i^* = 1 \tag{A.63a}$$

$$\pi_1^* = (1 - \sum_{j=1}^{2} m_j - \mu_1)\pi_1^* + \pi_8^* + \pi_{10}^* + \pi_{11}^* \tag{A.63b}$$

$$\pi_2^* = \mu_1 \pi_1^* \tag{A.63c}$$

$$\pi_3^* = (1 - \sum_{j=1}^{2} m_j)\pi_3^* + \pi_2^* + \pi_{12}^* \tag{A.63d}$$

$$\pi_4^* = m_2 \pi_3^* \tag{A.63e}$$

$$\pi_5^* = (1 - \sum_{j=1}^{2} m_j - \mu_1)\pi_5^* + \pi_4^* + \pi_9^* + \pi_{13}^* \tag{A.63f}$$

$$\pi_6^* = \mu_1 \pi_5^* \tag{A.63g}$$

$$\pi_7^* = (1 - \sum_{j=1}^{2} m_j)\pi_7^* + \pi_6^* + \pi_{14}^* \tag{A.63h}$$

$$\pi_8^* = m_1 \pi_7^* \tag{A.63i}$$

$$\pi_9^* = m_2 \pi_1^* \tag{A.63j}$$

$$\pi_{10}^* = m_1 \pi_5^* \tag{A.63k}$$

$$\pi_{11}^* = m_1 \pi_1^* \tag{A.63l}$$

$$\pi_{12}^* = m_1 \pi_3^* \tag{A.63m}$$

$$\pi_{13}^* = m_2 \pi_5^* \tag{A.63n}$$

$$\pi_{14}^* = m_2 \pi_7^*. \tag{A.63o}$$

From the above simultaneous linear equations, the elements of the stationary distribution are rewritten as follows:

$$
{}^{\mathrm{t}}\boldsymbol{\pi}^{*} = \begin{pmatrix} \pi_1^* \\ \mu_1\pi_1^* \\ \mu_1 m_2^{-1}\pi_1^* \\ \mu_1\pi_1 \\ (\mu_1+m_2)(\mu_1+m_1)^{-1}\pi_1^* \\ \mu_1(\mu_1+m_2)(\mu_1+m_1)^{-1}\pi_1^* \\ \mu_1(\mu_1+m_2)m_1^{-1}(\mu_1+m_1)^{-1}\pi_1^* \\ \mu_1(\mu_1+m_2)(\mu_1+m_1)^{-1}\pi_1^* \\ m_2\pi_1^* \\ m_1(\mu_1+m_2)(\mu_1+m_1)^{-1}\pi_1^* \\ m_1\pi_1^* \\ \mu_1 m_1 m_2^{-1}\pi_1^* \\ m_2(\mu_1+m_2)(\mu_1+m_1)^{-1}\pi_1^* \\ \mu_1 m_2(\mu_1+m_2)m_1^{-1}(\mu_1+m_1)^{-1}\pi_1^* \end{pmatrix}, \tag{A.64}
$$

where ${}^{\mathrm{t}}\boldsymbol{\pi}^*$ represents a transpose matrix of $\boldsymbol{\pi}^*$. As the sum of the elements of $\boldsymbol{\pi}^*$ should be one, one can find

$$
\pi_1^* = \frac{m_1 m_2 (m_1+\mu_1)}{A}, \tag{A.65}
$$

where

$$
\begin{aligned} A \quad = \quad & m_1 m_2 (m_1+\mu_1)(m_1+m_2+2\mu_1+1) + m_1 m_2 (m_2+\mu_1)(m_1+m_2+2\mu_1+1) \\ & + m_1\mu_1 (m_1+1)(m_1+\mu_1) + m_2\mu_1 (m_2+1)(m_2+\mu_1). \end{aligned} \tag{A.66}
$$

Now, we analytically obtain the stationary distribution $\boldsymbol{\pi}^*$. The expected cumulative efficiency of detoxification from the beginning of cultivation to time step $s+1$, $\Phi(s+1;\boldsymbol{m},\boldsymbol{\mu})$, is given by:

$$
\Phi(s+1;\boldsymbol{m},\boldsymbol{\mu}) = \Phi(s;\boldsymbol{m},\boldsymbol{\mu}) + \sum_{i=1}^{14} C\phi_i\pi_i(s+1;\boldsymbol{m},\boldsymbol{\mu}), \tag{A.67}
$$

where $C$ is a positive constant to change the time scale of $\phi_i$ into a discrete time step. The second term of Eq (A.67) represents the expected detoxification efficiency at time step $s+1$. The optimization problem is to find the values of $m_1$ and $m_2$ that maximize the expected cumulative efficiency given time step $s$, which is solved by Dynamic Programming. However, in the limit of $s \to \infty$, $\boldsymbol{\pi}(s,\boldsymbol{m};\boldsymbol{\mu}) \to \boldsymbol{\pi}^*(\boldsymbol{m};\boldsymbol{\mu})$ due to the ergodicity of the Markov chain. Once we reach the stationary distribution $\boldsymbol{\pi}^*$, the second term of Eq (A.67) becomes $C\hat{\Phi}(\boldsymbol{m};\boldsymbol{\mu})$, which is independent on time step $s$ (see Eq (11)).

Suppose that the probability distribution is enough close to $\boldsymbol{\pi}^*$ since time step $s^*$. At time step $s > s^*$,

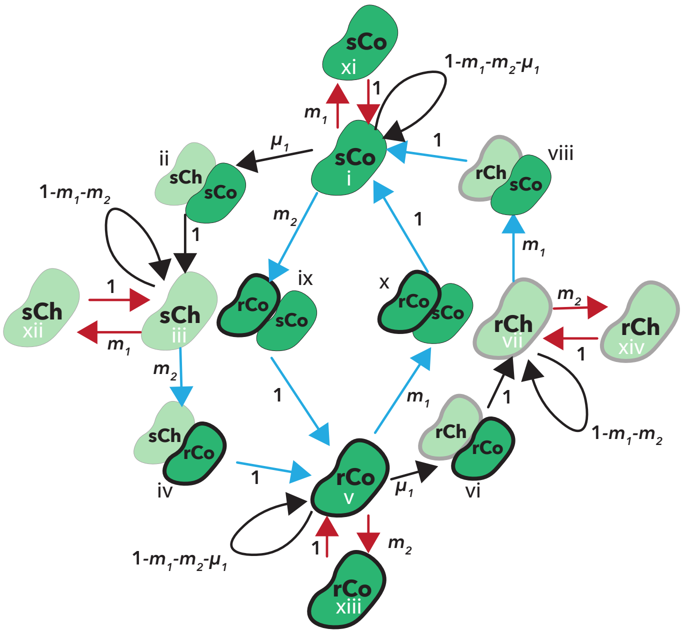


Figure A.9: The diagram of state transitions when mutation never occurs in the level of resistance

The diagram of the state transitions when $\mu_2 = 0$. In this example, there exist 14 states; mono-cultures of one of four strategies, transient states caused by the invasion of cheaters or by the introduction of cooperators with different resistance levels (sky blue arrows), and the unnecessary introduction of cooperators which currently exist or cannot invade the population (red arrows). Each value along the arrows represents the corresponding transition probability. The numbers in the roman numerals correspond to state $i$; $i = 1$ represents the mono-culture of sCo. The optimal introduction rates of the cooperators in this model are shown in the bottom panel of Fig. 5.

the cumulative expected detoxification efficiency is approximately:

$$\Phi\left(s;\boldsymbol{m},\boldsymbol{\mu}\right) \approx \left(s - s^* + 1\right) C\hat{\Phi}\left(\boldsymbol{m};\boldsymbol{\mu}\right) + \Phi\left(s^* - 1;\boldsymbol{m},\boldsymbol{\mu}\right). \tag{A.68}$$

At large $s$, the second term of Eq (A.68) is small relative to its first term. The optimization problem for the cumulative expected detoxification efficiency then simplifies to maximizing Eq (11). In other words, for a large time step, the optimal values of $m_1$ and $m_2$ converge to those which maximize Eq (11) as shown in Fig. 5B.

Next, we relax the three assumptions in Box 1. If a mutation occurs in the transient states, or if a mutation and the intentional introduction of cooperators occur at the same time (relaxing assumption (ii)), the Markov chain does not lose ergodicity (Fig. A.10 top left). Although relaxing this assumption allows the population to hold three strategies at once, there exist at most two strategies in the population at an equilibrium state (see Appendix 3 and Appendix 4).

If cooperators cannot invade a population of cooperators with different resistance level (relaxing assumption (iii)), (a) resident cooperators remain in the population but detoxification efficiency is not optimized in that state, or (b) resident cooperators also go extinct because they cannot persist at $(\alpha, T_{\text{in}})$ which maximizes detoxification efficiency in a mono-culture of invader cooperators (Fig. A.10 top right). In both cases, however, the Markov chains are still ergodic. In case (a), resident cooperators are excluded by cheaters that have the same level of resistance regardless of the values of $(\alpha, T_{\text{in}})$. In case (b), on the other hand, sensitive or resistant cooperators can be introduced with probability $m_1$ and $m_2$ when the population size is 0. Indeed, if all microbes go extinct, the Markov chains are always ergodic because the state can transit to a mono-culture of either type of cooperator from this state. Therefore, the optimal $\boldsymbol{m}$ is found in the same way as in the main text.

If sCo and/or rCo cannot exclude the cheaters that differ in the level of resistance (relaxing assumption (iv)), the Markov chains can be non-ergodic. The Markov chain is not ergodic if the initial condition is a mono-culture of one type of cooperators where the population state never goes back once it has changed. When the initial condition is a mono-culture of sCo and sCo cannot exclude rCh, for example, the Markov chain is not ergodic if the state transition cannot occur from a mono-culture of rCo or the coexistence of rCo with sCh to a mono-culture or sCo (Fig. A.10 bottom left and right). Note that although cooperators cannot invade the population where the other type of cooperators and the cheaters that differ in the level of resistance coexist, the manual introduction of cooperators can collapse this coexistence by changing $(\alpha, T_{\text{in}})$. In addition, the values of $(\alpha, T_{\text{in}})$ in a mono-culture of sCo should be the same as in the coexistence of sCo with rCh; otherwise, the initial condition is transient and ergodicity is lost.

Now we begin the analysis of non-ergodic Markov chains. When sensitive (and/or resistant) cooperators cannot exclude cheaters with different resistance level by changing the culture conditions, the Markov chains are not always ergodic. Indeed, if the initial condition is a mono-culture of cooperators that cannot exclude cheaters with different level of resistance, and if there is no state transition from another state to the mono-culture of this type of cooperator, the Markov chain is non-ergodic. In this case, the set of states is divided into two subsets $S_1$ and $S_2$. Subset $S_1$ is composed of the equilibrium states where there exist only cooperators that cannot exclude cheaters with different levels of resistance (the values of two parameters, $\alpha$ and $T_{\text{in}}$ are different at each state), and the transient states from these equilibrium states to other equilibrium states in subset $S_1$. Subset $S_2$, on the other hand, includes the rest of the states within the state space, and the Markov chain is ergodic within $S_2$. In such cases, the expected cumulative efficiency of detoxification is recursively represented as below:

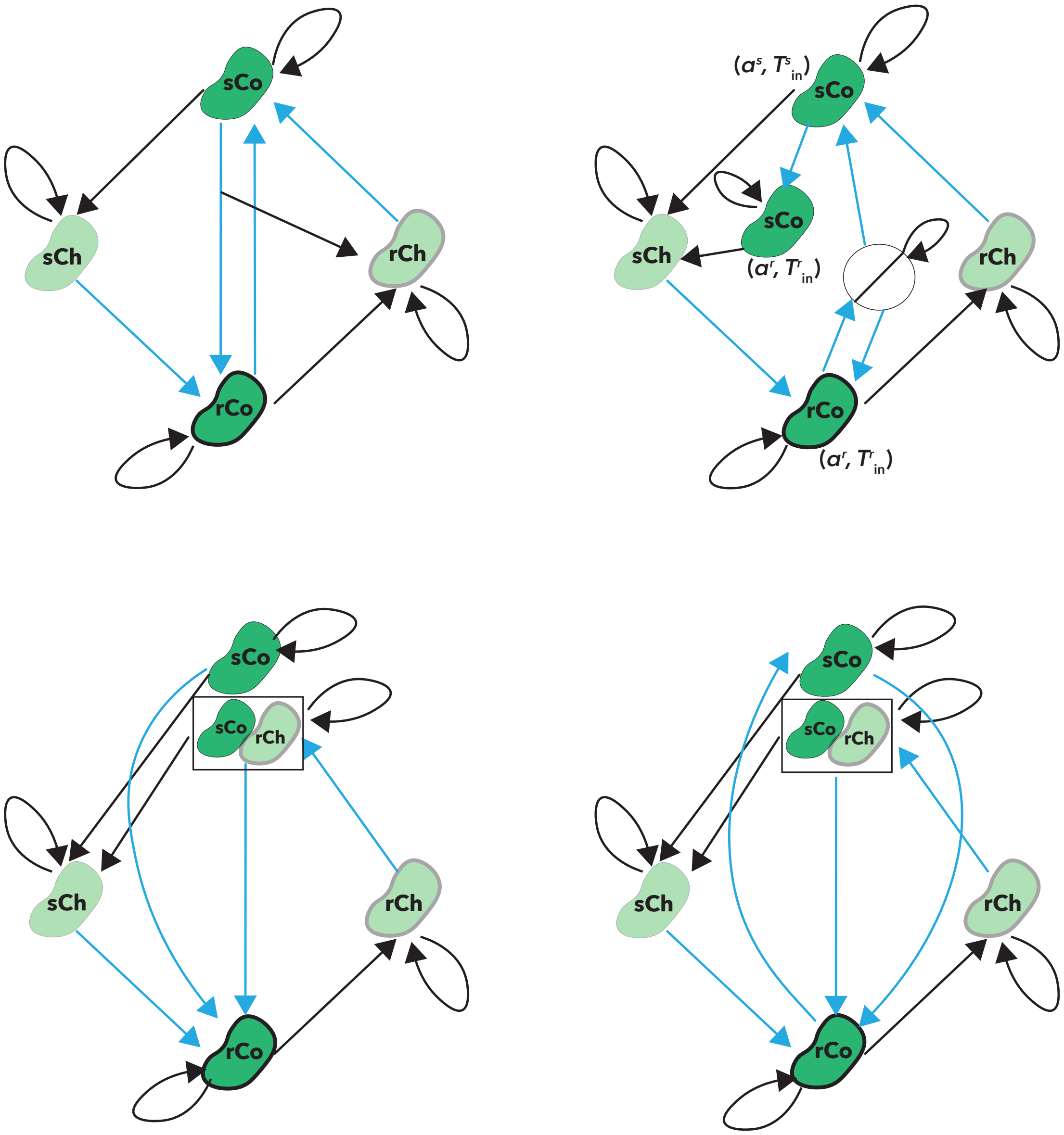


Figure A.10: Diagrams of state transitions

In each panel, the transient states and the states where the manual introduction of cooperators decreases the efficiency of detoxification is ignored. In addition, it is assumed that $\mu_2 = 0$ to reduce the number of nodes. The black and sky blue lines represent the state transitions caused by mutation and manual introduction, respectively. Top left: when rCo is introduced into a population of sCo, rCo can mutate into rCh, which can add the node to rCh. Top right: if rCo cannot invade sCo, sCo persists in the population. However, the values of $(\alpha, T_{\text{in}})$ change and the efficiency of detoxification differs before the invasion. If sCo cannot invade rCo and rCo goes extinct where the efficiency of detoxification is maximized for a population of sCo, the population state transits from rCo to the white, slashed circle where all microbes go extinct. If sCo or rCo are introduced, however, they can grow in the chemostat. Bottom left: if sCo cannot exclude rCh and sCo cannot invade a population of rCo, the Markov chain is not ergodic, when the initial condition is a mono-culture of rCo. Notice, however, that the Markov chain can be regraded as ergodic by ignoring the state of sCo when the initial condition is rCo. Bottom right: Although sCo cannot exclude rCh, the Markov chain can be ergodic when sCo can exclude rCo.

$$
\begin{aligned}
\Phi(s+1,\boldsymbol{m};\boldsymbol{\mu}) &= \Phi(s,\boldsymbol{m};\boldsymbol{\mu}) \\
&+ \prod_{i=0}^{s+1}\left\{1-p_{12}\left(i,\boldsymbol{m};\boldsymbol{\mu}\right)\right\}\sum_{k\in S_1} C\phi_k\pi_k\left(s+1,\boldsymbol{m};\boldsymbol{\mu}\right) \\
&+ \left[1-\prod_{i=0}^{s+1}\left\{1-p_{12}\left(i,\boldsymbol{m};\boldsymbol{\mu}\right)\right\}\right]\sum_{k\in S_2} C\phi_k\pi_k\left(s+1,\boldsymbol{m};\boldsymbol{\mu}\right),
\end{aligned}
\tag{A.69}
$$

where $p_{12}(i)$ represents the state transition probability from a state in $S_1$ to another in $S_2$ at $s = i$. Then, the second and third terms of the above equation represent the expected detoxification efficiency when the state of the population is in subset $S_1$ or $S_2$, respectively.

To calculate the state distribution in this case, we separate the probability distribution vector into two vectors, $\boldsymbol{\pi}_i = (\pi_j)$ where $j \in S_i$ for $i = 1, 2$. Then, each vector at time step $s + 1$ is calculated as below:

$$
\boldsymbol{\pi}_1\left(s+1,\boldsymbol{m};\boldsymbol{\mu}\right) = \boldsymbol{\pi}_1\left(s,\boldsymbol{m};\boldsymbol{\mu}\right)P^{11}\left(\boldsymbol{m};\boldsymbol{\mu}\right), \tag{A.70a}
$$

$$
\boldsymbol{\pi}_2\left(s+1,\boldsymbol{m};\boldsymbol{\mu}\right) = \boldsymbol{\pi}_1\left(s,\boldsymbol{m};\boldsymbol{\mu}\right)P^{21}\left(\boldsymbol{m};\boldsymbol{\mu}\right)+\boldsymbol{\pi}_2\left(s,\boldsymbol{m};\boldsymbol{\mu}\right)P^{22}\left(\boldsymbol{m};\boldsymbol{\mu}\right), \tag{A.70b}
$$

where matrix $P^{ij}$ represents the transition probability from a state in $S_i$ to another in $S_j$. Note that $p_{12}\left(i,\boldsymbol{m};\boldsymbol{\mu}\right)$ is given by the sum of all elements of $\boldsymbol{\pi}_1\left(i,\boldsymbol{m};\boldsymbol{\mu}\right)P^{21}\left(\boldsymbol{m};\boldsymbol{\mu}\right)$.

As there is no state transition from a state in $S_2$ to another in $S_1$, $\boldsymbol{\pi}_1\left(s,\boldsymbol{m};\boldsymbol{\mu}\right) \to \mathbf{0}$ in the limit of $s \to \infty$. Once $\boldsymbol{\pi}_1\left(s,\boldsymbol{m};\boldsymbol{\mu}\right)P^{21}\left(\boldsymbol{m};\boldsymbol{\mu}\right)$ becomes small enough, the state distribution $\boldsymbol{\pi}_2$ is approximated as:

$$
\boldsymbol{\pi}_2\left(s+1,\boldsymbol{m};\boldsymbol{\mu}\right) \approx \boldsymbol{\pi}_2\left(s,\boldsymbol{m};\boldsymbol{\mu}\right)P^{22}\left(\boldsymbol{m};\boldsymbol{\mu}\right). \tag{A.71}
$$

In the limit of $s \to \infty$, therefore, $\boldsymbol{\pi}_2$ converges to a unique stationary distribution $\boldsymbol{\pi}_2^*$ as a Markov chain given by subset $S_2$ is ergodic. In addition, $\prod_{i=0}^{s}\left\{1-p_{12}\left(i,\boldsymbol{m};\boldsymbol{\mu}\right)\right\}$ in Eq(A.69) converges to zero with large $s$. Therefore, Eq (A.69) is simplified in the limit of $s \to \infty$ as below:

$$
\Phi(s+1,\boldsymbol{m};\boldsymbol{\mu}) \approx \Phi(s,\boldsymbol{m};\boldsymbol{\mu}) + \sum_{k\in S_2} C\phi_k\pi_k^*\left(\boldsymbol{m};\boldsymbol{\mu}\right). \tag{A.72}
$$

Thus, the optimal $\boldsymbol{m}$ is approximately obtained by maximizing $\sum_{k\in S_2}\phi_k\pi_k^*\left(\boldsymbol{m};\boldsymbol{\mu}\right)$.

Let us show a simple example of a non-ergodic Markov chain. Here, we assume that sCo cannot exclude rCh at any $\left(\alpha, T_{\text{in}}\right)$. In addition, it is assumed that neither cooperator can invade the population of the other type of cooperator, and that no state exists where all cells go extinct. In this case, the Markov chain is ergodic if the initial state is a mono-culture of sCo. For simplicity, we ignore transient states and assume that $\mu_2 = 0$ as in the main text (Fig. A.11 left). At more than $1,000$ time steps, the

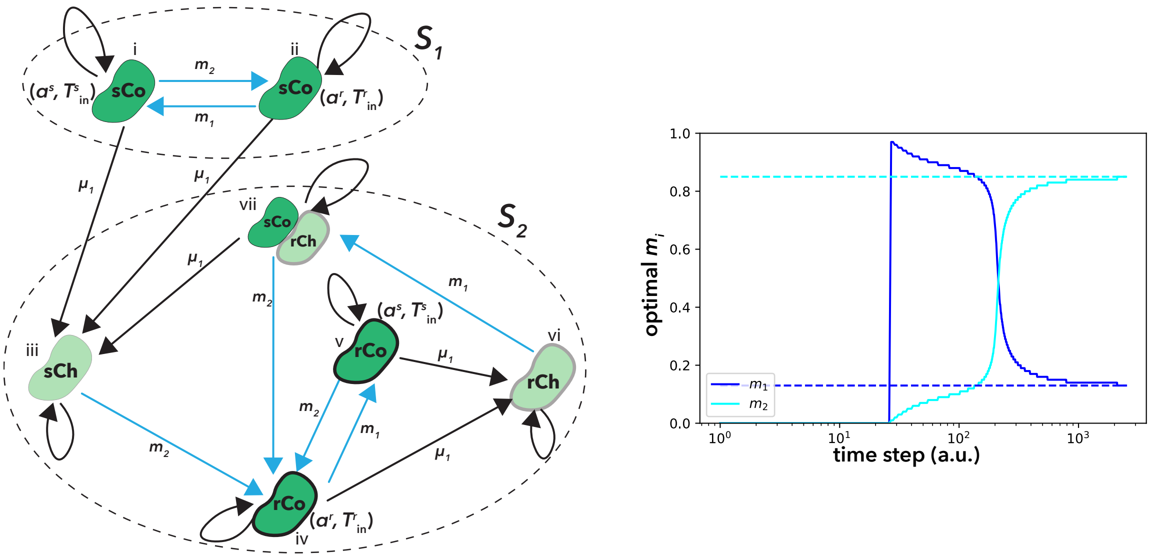


Figure A.11: An example when a Markov chain is not ergodic

Left: the state transition diagram of a non-ergodic Markov chain is shown. In this example, it is assumed that sCo cannot exclude rCh, that $\mu_2 = 0$, and that both types of cooperators cannot invade each other. Therefore, when one type of cooperators is introduced to the population of the other type of cooperators, the resident cooperators remain in the population but the values of $\alpha$ and $T_{\text{in}}$ are changed. To reduce the number of states, the transient states are ignored. Notice that there is no state transition from subset $S_2$ to $S_1$. In addition, the Markov chain within subset $S_2$ is ergodic. Each number corresponds to the index of each state. Right: the optimal $\boldsymbol{m}$ (the solid lines) when the simulation finishes at each time step is shown when $\mu_1 = 0.01$ in the left panel. The dashed lines represent $\boldsymbol{m}$ which maximizes $\sum_{k\in S_2} \phi_k \pi_k^* (\boldsymbol{m}; \boldsymbol{\mu})$. The other parameter values are $\boldsymbol{\phi} = \{0.4, 0.2, 0, 0.3, 0.15, 0, 0.2\}$.

optimal $\boldsymbol{m}$ converges to the values where $\sum_{k\in S_2} \phi_k \pi_k^* (\boldsymbol{m}; \boldsymbol{\mu})$ is maximized (Fig. A.11 right).
